# Anatomically distributed neural representations of instincts in the hypothalamus

**DOI:** 10.1101/2023.11.21.568163

**Authors:** Stefanos Stagkourakis, Giada Spigolon, Markus Marks, Michael Feyder, Joseph Kim, Pietro Perona, Marius Pachitariu, David J. Anderson

## Abstract

Artificial activation of anatomically localized, genetically defined hypothalamic neuron populations is known to trigger distinct innate behaviors, suggesting a hypothalamic nucleus-centered organization of behavior control. To assess whether the encoding of behavior is similarly anatomically confined, we performed simultaneous neuron recordings across twenty hypothalamic regions in freely moving animals. Here we show that distinct but anatomically distributed neuron ensembles encode the social and fear behavior classes, primarily through mixed selectivity. While behavior class-encoding ensembles were spatially distributed, individual ensembles exhibited strong localization bias. Encoding models identified that behavior actions, but not motion-related variables, explained a large fraction of hypothalamic neuron activity variance. These results identify unexpected complexity in the hypothalamic encoding of instincts and provide a foundation for understanding the role of distributed neural representations in the expression of behaviors driven by hardwired circuits.

## Introduction

Brains evolved to create a series of abstractions that represent objects or other sensory cues in an organism’s environment, assess their value, predict impending changes in their features, and transform those signals into adaptive behavioral responses. To deconstruct this complex process, two main experimental strategies have been taken to either: 1) record neural activity during behavior, or 2) manipulate neural activity and observe changes in behavior. The first strategy starts with measurements of neuronal activity and correlates those data with manipulations and measurements of sensory features or task variables^1–5^. The behavioral readouts typically involve artificial tasks that require intensive training and are highly reproducible across trials. In mammals, this strategy has been employed to study the cortex, hippocampus, or other evolutionarily recent areas^6–10^, and their influence on cognitive functions.

The second strategy starts with surgical, electrical, neurochemical, or genetically-based perturbation experiments to identify brain regions and cell types whose activation, inhibition, or destruction can promote or suppress particular behaviors. These manipulations are targeted to pre-selected areas based on hypotheses, assumptions, or prior knowledge. This strategy has dominated the study of internal states, motivation, reward, and emotion^11–15^. The behavioral readouts typically comprise innate behaviors or simple forms of conditioning based on such instincts. The study of such functions has typically focused on the so-called “limbic system,” including evolutionarily ancient regions such as the extended amygdala^16,17^, hypothalamus^18,19^, and midbrain^20,21^.

The two above-mentioned strategies can yield fundamentally different views of the mechanisms underlying brain functions of interest. Although many studies in practice employ both strategies sequentially, prevailing views in a field are often strongly shaped by the approach that was initially undertaken^22–24^. The “recording first” strategy has led to an emphasis on high-dimensional population codes, neural dynamics, and manifolds in cortico-hippocampal state space. Nevertheless, the field is increasingly employing perturbation experiments to test hypotheses based on observational studies. In contrast, the “perturbation first” strategy has led to an emphasis in the hypothalamus field on hard-wired neural circuits that causally control specific innate behaviors in a manner that is highly deterministic and low-dimensional^25–27^. However, large-scale recording experiments in this field have been relatively rare.

It remains unclear, therefore, whether these different views of how cortical and limbic systems control behavior reflect true biological differences, or rather differences in the evolution of these fields according to the initial experimental strategies used at their inception. To help resolve this discrepancy, here we have performed large-scale recordings from neural populations located along the rostrocaudal axis of the medial hypothalamus, using custom-designed silicon probes in freely moving mice engaged in solitary, social, or predator-defense behaviors. Our results reveal that the neural coding of such instinctive behaviors is, surprisingly, more distributed, dynamic, and high-dimensional than would have been expected from perturbation experiments. These results invite a re-evaluation of views of neural coding principles in cortical vs. limbic areas and highlight the importance of integrating results from observational and perturbational studies to arrive at a comprehensive understanding of brain function.

## Results

### Recording single neurons across multiple sites of the hypothalamus during behavior

We sought to acquire synchronized large-scale electrophysiological and video recording data from freely moving mice engaging in diverse innate behaviors. To record the activity of hundreds of neurons simultaneously throughout the medial hypothalamus, we used a custom-fabricated silicon probe (NeuroNexus, Inc.) that allows chronic extracellular electrophysiological recordings (with single-unit resolution) from multiple nuclei (Figure 1A). The eight-shank 256-channel silicon probe (Figure 1B) was designed to provide simultaneous access to twenty hypothalamic subregions (nuclei or subnuclei), which are spatially distributed across the rostrocaudal extent of the hypothalamus (Figure 1E). The probe covers ∼2.3mm of brain tissue (Figure 1A, lower), and we recorded ∼400 units per animal (n=6 animals) for a total of 2,347 units.

**Figure 1.**
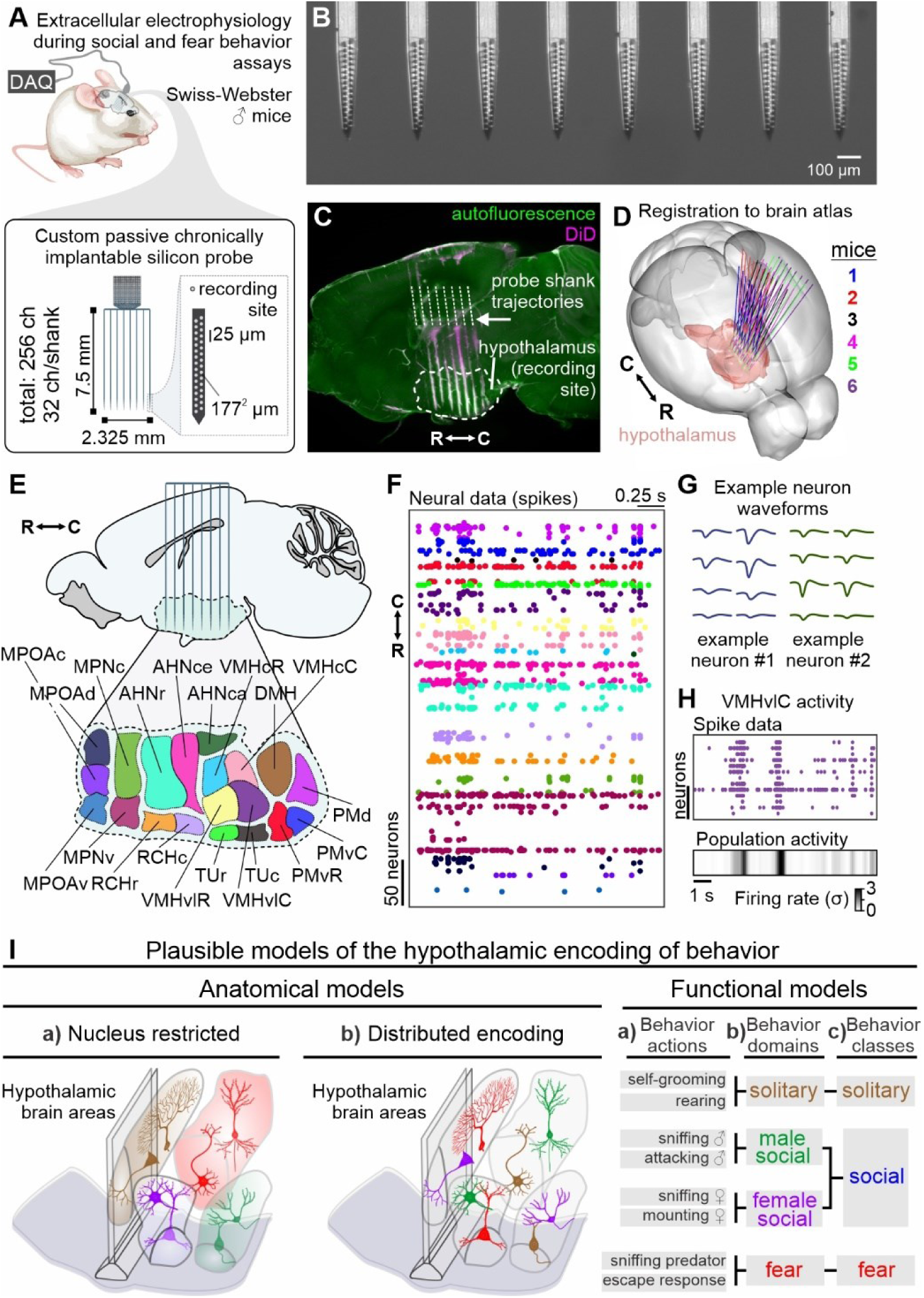
Approach for single neuron recordings across the hypothalamus using custom extracellular electrophysiology silicon probes in freely moving mice. (A) Top - outbred Swiss Webster male mice were chronically implanted with a custom silicon probe. Bottom - design, and specifics of the eight-shank 256-channel custom silicon probe manufactured by NeuroNexus. (B) Stereoscope-acquired image of the shank-tips of the probe containing the microelectrodes. (C) Post-hoc identification of probe shanks and electrode placement through volumetric light-sheet imaging. (D) Registration of the probes to the Allen Common Coordinate Framework across all recorded mice. (E) Target coordinates include twenty, individually colored subregions of the hypothalamus. (F) Following electrode mapping and spike sorting, single neurons were assigned to their respective locations / hypothalamic areas. (G) Example waveforms of two recorded neurons. (H) Example spike data across thirteen neurons of VMHvl (top), and illustration of their average activity (bottom). (I) Spatial and functional plausible models, combination of which could explain the hypothalamic encoding of behavior. Abbreviations: DAQ: data acquisition system, MPOAv: medial preoptic area – ventral subdivision. MPOAc: medial preoptic area – central subdivision. MPOAd: medial preoptic area – dorsal subdivision. MPNv: medial preoptic nucleus – ventral subdivision. MPNc: medial preoptic nucleus – central subdivision. RCHr: retrochiasmatic area – rostral subdivision. RCHc: retrochiasmatic area – caudal subdivision. AHNr: anterior hypothalamic nucleus – rostral subdivision. AHNce: anterior hypothalamic nucleus – central subdivision. AHNca: anterior hypothalamic nucleus – caudal subdivision. TUr: tuberal nucleus – rostral subdivision. TUc: tuberal nucleus – caudal subdivision. VMHvlR: ventromedial hypothalamic nucleus ventrolateral segment – rostral subdivision. VMHvlC: ventromedial hypothalamic nucleus ventrolateral segment – caudal subdivision. VMHcR: ventromedial hypothalamic nucleus central segment – rostral subdivision. VMHcC: ventromedial hypothalamic nucleus central segment – caudal subdivision. DMH: dorsomedial hypothalamic nucleus. PMvR: ventral premammillary nucleus – rostral subdivision. PMvC: ventral premammillary nucleus – caudal subdivision. PMd: dorsal premammillary nucleus. See also Figures S1 and S2.

To map shank trajectories, the back side of the probe shanks was coated with a red lipophilic dye that could be traced post-hoc. Intact iDISCO-cleared mouse brains were volumetrically imaged using light-sheet microscopy^28^, and probe coordinates were mapped to the Allen Common Coordinate Framework^29^ (Figure 1C and 1D). A unique electrode map was generated for each mouse by manually mapping electrodes to computationally sectioned sagittal sections of each mouse brain (Figures S1A-S1B). A schematic map of the areas recorded in each mouse is shown in Figure 1E. Based on the electrode maps, neurons were allocated to specific hypothalamic brain areas (Figure 1F, single-neuron activity colored according to the location of each neuron). Spike-sorting was performed using Kilosort^30^, and basic analysis of neuron waveforms (Figure 1G), and quality control of recordings was performed using pre-existing pipelines (see also STAR Methods section: *in vivo* electrophysiology). Neural data were analyzed in relation to behavior either a) with single neuron resolution (Figure 1H – upper part), or b) by averaging the activity of cells per nucleus (Figure 1H – lower part). This experimental design enabled us to test whether the encoding of behavior in the hypothalamus conforms to anatomical nuclear boundaries and examine the extent to which neurons from the same nucleus are highly correlated or have distinct activity patterns across diverse behaviors. (Figure 1I).

To investigate the representation of behavior in the hypothalamus, we grouped behaviors into three hierarchical levels (Figure 1I, right). At the highest level, we defined three broad behavioral categories which we refer to as “classes“: the solitary class includes actions such as self-grooming, and rearing. The social class includes behaviors such as sniffing a male or female conspecific or attacking a male conspecific. The fear class includes behaviors such as sniffing a predator or exhibiting an escape response. At the second level, we distinguished two “domains” within the “social” class: intermale and male-female social behaviors (Figure 1I, right; green and violet, respectively). Finally, at the third and lowest level, we annotated 43 distinct actions among the different classes and domains (Figure 3B).

To ensure that the behaviors of interest would be expressed during the neural data recording sessions, animals (male Swiss-Webster mice) were pre-screened for robust performance in the relevant behavioral assays (Figure 2A, a) solitary, b) social, and c) fear). This pre-screening step reinforced the expression of social behaviors^11,12^, and the display of fear behaviors during the recording sessions (Figure S1C-S1I). Following the probe implantation, animals were allowed to recover and were habituated to the experimental setup in the absence of the behavioral assays (Figure 2B).

**Figure 2.**
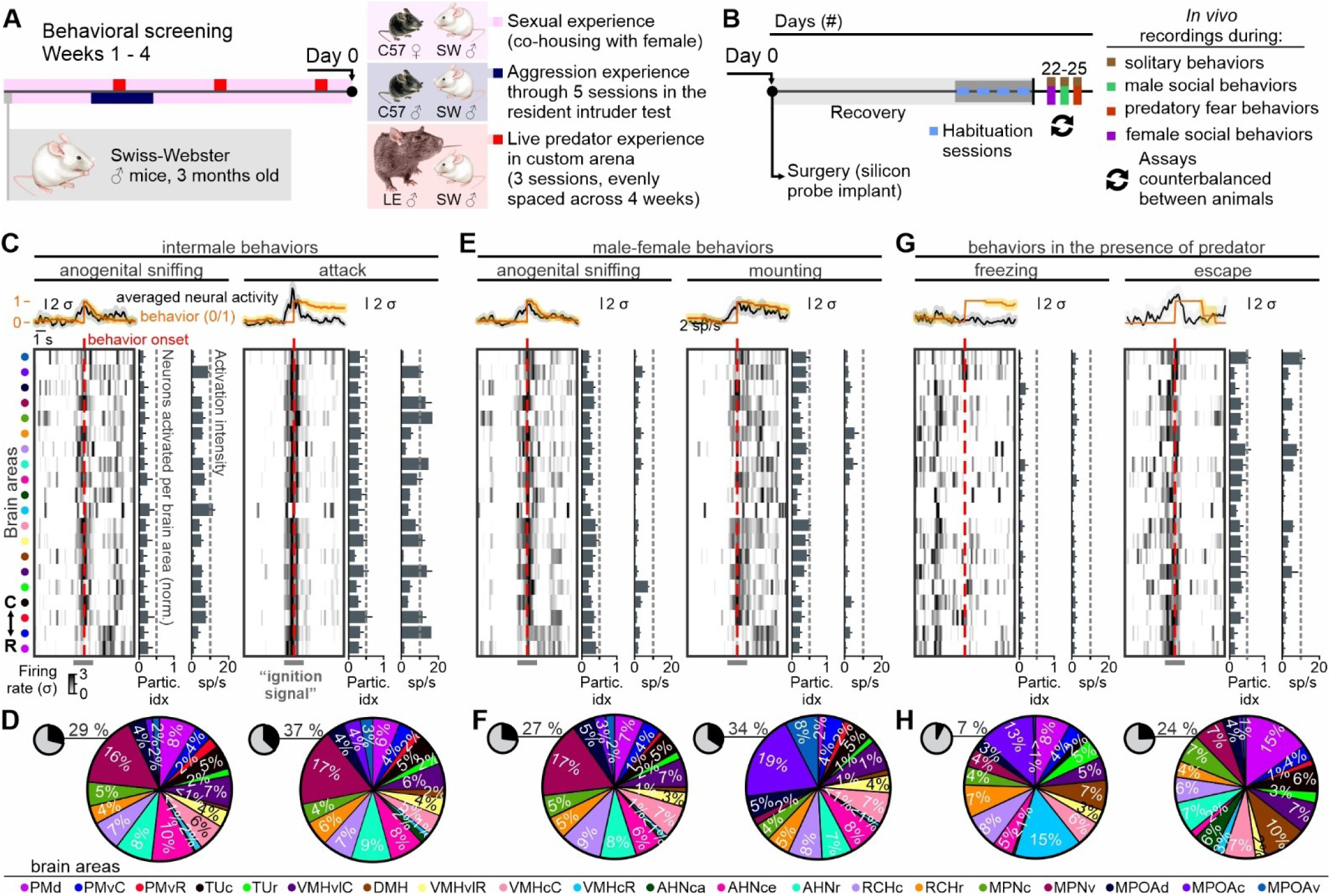
Neural ignition signals dominate the peri-onset interval of all instinctual behaviors, apart from freezing to the presence of a predator. (A) Behavioral screening of outbred Swiss Webster mice was used to identify individual mice with prominent behavioral expressions in all of the three behavior paradigms of interest used to assess mating with a female conspecific, aggression against a male conspecific and fear behaviors in the presence of a predator (Long-Evans [LE] rat). Only mice that exhibited appetitive and consummatory behaviors across the three behavior domains (i.e., mating, aggression, and fear), across all behavior screening sessions were selected for chronic silicon probe implant surgery. (B) Schematic illustrating the timeline following implantation of the silicon probe. Animals were allowed to fully recover for two weeks in their home cage and were given access to a running wheel. Four habituation sessions were performed (one daily), 4 days prior to the start of the neural recordings and behavior experiments. Habitation sessions included exposure to the exact experimental conditions, in the absence of the stimulus (conspecific, or predator). (C) Average behavior signal and mean neural activity at the peri-onset interval of an appetitive and a consummatory intermale social behavior (top). Average neural activity per nucleus (heatmap), participation index (% of neurons activated above 2 SD from baseline activity), and average firing rate of the activated neurons (bottom, n = 6 mice, neural data averaged across three consecutive behavior instances per mouse). (D) Small black and gray pie plot: Black – total % of neurons activated, Gray: total % of neurons that did not raise their activity above 2σ. Larger colored pie plot illustrates the spatial coordinates of the neurons that became active (raised their activity >2σ), during male anogenital sniffing (left) and intermale attack (right, n = 6 mice, neural data averaged across three consecutive behavior instances). (E and F) Same as panels (C) and (D) for an appetitive and a consummatory male-female social behavior. (G and H) Same as panels (C) and (D) for two predatory fear behaviors. In box-and-whisker plots, center lines indicate medians, box edges represent the interquartile range, and whiskers extend to the minimal and maximal values. See also Figures S3, S4, and S5. Abbreviations: Partic. Idx.: participation index, sp/s: Firing rate (spikes / sec). Abbreviations of brain areas are described on the legend of Figure 1.

**Figure 3.**
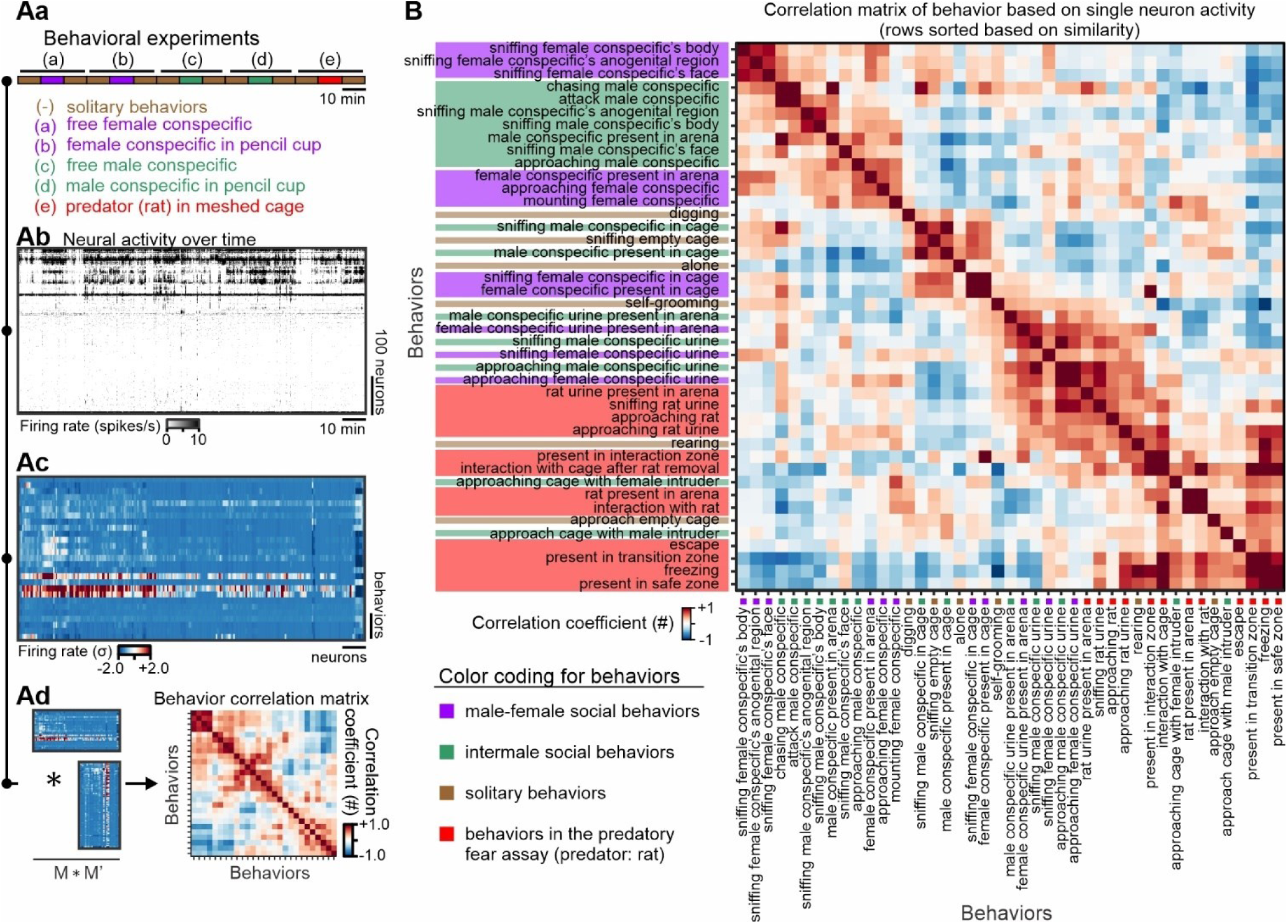
Non-overlapping encoding of social and fear behavior classes by hypothalamic neurons. (A) Analysis design to generate a behaviors correlation matrix. (Aa) Experimental conditions. (Ab) Single-unit neural activity during the behavioral paradigms shown in Aa. (Ac) Z-scored activity of all recorded neurons across solitary-, social-, and fear-behaviors (data from an example mouse). (Ad) Multiplication of the matrix shown in ac with its transposed version, results to the behavior covariance matrix, which is then transformed to a behavior correlation matrix. (B) Correlation matrix of behavior (average across n = 6 mice). Correlation matrices of behavior of the individual mice can be found in Figure S7.

As previous work has shown that neural firing rates often correlate with motion-related variables in some brain areas^31–33^, we assessed this question in our mesoscale hypothalamic recordings. Self-motion variables and distance between animals did not correlate with mean single unit firing rates, either during whole recording sessions (Figures S2A and S2Ba-S2Bc), or during specific behavior actions (Figure S2Bd). Partial least squares (PLS) regression provided further evidence that single-unit activity also does not correlate with motion variables (Figure S2C, note a near zero R^2^ between PLS components of neural data and of motion-related variables in Figure S2Cd). However, the PLS regression revealed that single-unit activity was correlated with consummatory (Figure S2D, note a high R^2^ between PLS components of neural data and of consummatory behaviors in Figure S2Dd), and, to a lesser extent, with appetitive (Figure S2E, note a medium range R^2^ between PLS components of neural data and of appetitive behaviors in Figure S2Ed) social and fear behaviors.

### Spatially distributed and behavior non-specific neural “ignition signals”

Next, we sought to address whether activity during specific appetitive and consummatory behaviors, including intermale social behaviors (Figure 2C and 2D), male-female social behaviors (Figure 2E and 2F), and predatory defensive behaviors (Figure 2G and 2H), is restricted anatomically or rather spatially distributed, based on averaging units recorded from each of the twenty nuclei. Throughout the entire hypothalamus, we observed a synchronous burst of activity near the onset of all behaviors except for predator-induced freezing (Figure 2C-2G). This spiking burst that occurred at all behavior onsets following the introduction of an intruder, appeared consistently across behavioral episodes (Figure S2F and S2G). It had an onset of -200 msec prior to the initiation of behavior (Figure S2H and S2Ia), an average duration of 1.5 sec (Figure S2Ib), and always ended before the termination of the behavior (Figure S2Ic). This response, which we refer to here as an “ignition signal,” encompassed ∼25-35% of recorded units across the entire hypothalamus, except for freezing behavior which engaged only 7% of units (Figure 2D-H, gray/black pie charts). The ignition signal was observed across multiple episodes of a given behavior directed towards the same intruder or predator (Figure S2F), indicating that it did not simply represent a novelty or neophobic response. However, the activity levels and proportion of the neurons recruited across behaviors markedly differed across nuclei (Figure 2C, 2E, and 2G – right bar plots). Each nucleus was differentially activated during behaviors of interest (Figure 2D, 2F, and 2H). Together, these findings indicate that ignition signals, while globally distributed, exhibit quantitative biases in their anatomic distribution that likely reflect behavior-specific responses in different hypothalamic nuclei (Figure 2D, 2F, and 2H).

The near-uniform expression of ignition signals likely accounts for the lack of statistical significance in decoding behaviors from nucleus-averaged neural activity (SVM decoding models, Figure S2J). An analysis of activity at single-unit resolution – discussed in the next section, however, could help to address whether behavior-specific modes of activity exist at the level of individual neurons.

### Single neuron activity primarily encodes distinct behavior classes

To investigate the representation of behavior by single neurons in the hypothalamus, we annotated 43 behavioral actions among the different classes and domains (functional models discussed in Figure 1I, right). Using these annotations, the first approach we used was to compute the single neuron activity-based behavior correlation matrix (Figure 3Aa-3Ad). Using rastermap^34^, we sorted the row and column indices of the behavior correlation matrix to maximize similarity between consecutive rows and columns, which allowed us to evaluate the differences in neural representations across behaviors visually (Figure 3B). Following averaging of the behavior correlation matrix across six mice, we observed a prominent segregation between social and predatory-defensive behavioral classes (note the separation of red- [“fear“] vs. green- and purple-tagged [social] behaviors in Figure 3B). This effect was also prominent in the behavior correlation matrices of individual mice (Figure S3Aa-S3Af, with the notable exception of mouse #4, whose behavior correlation matrix is shown in Figure S3Ad). Overall, this analysis revealed that social and fear behavioral classes can be broadly segregated based on neuron population activity. Male- and female-directed social behaviors were also generally separated, but not as prominently. Related individual actions had correlated neural patterns to some extent within domains (Figure 3B, sorting of behavior actions reflects similarity in their encoding).

The results of the behavior correlation matrix suggested that social and fear behaviors are encoded differently by hypothalamic neurons. However, this population-level analysis cannot distinguish whether individual behaviors are encoded by the same set of neurons that exhibit behavior class-specific activity patterns or whether distinct groups of neurons encode distinct behavior classes. To address this question, we next used a more targeted method, focusing on only two representative behavioral actions from each of the four behavioral domains of interest (a total of eight behavioral actions, Figure 4A). This approach could 1) confirm or disprove the segregation of social vs. fear behaviors, and 2) if the former is the case, indicate whether this segregation reflects different neuron groups active in the two behavioral classes, or whether it is driven by different patterns of activity exhibited by the same neurons.

**Figure 4.**
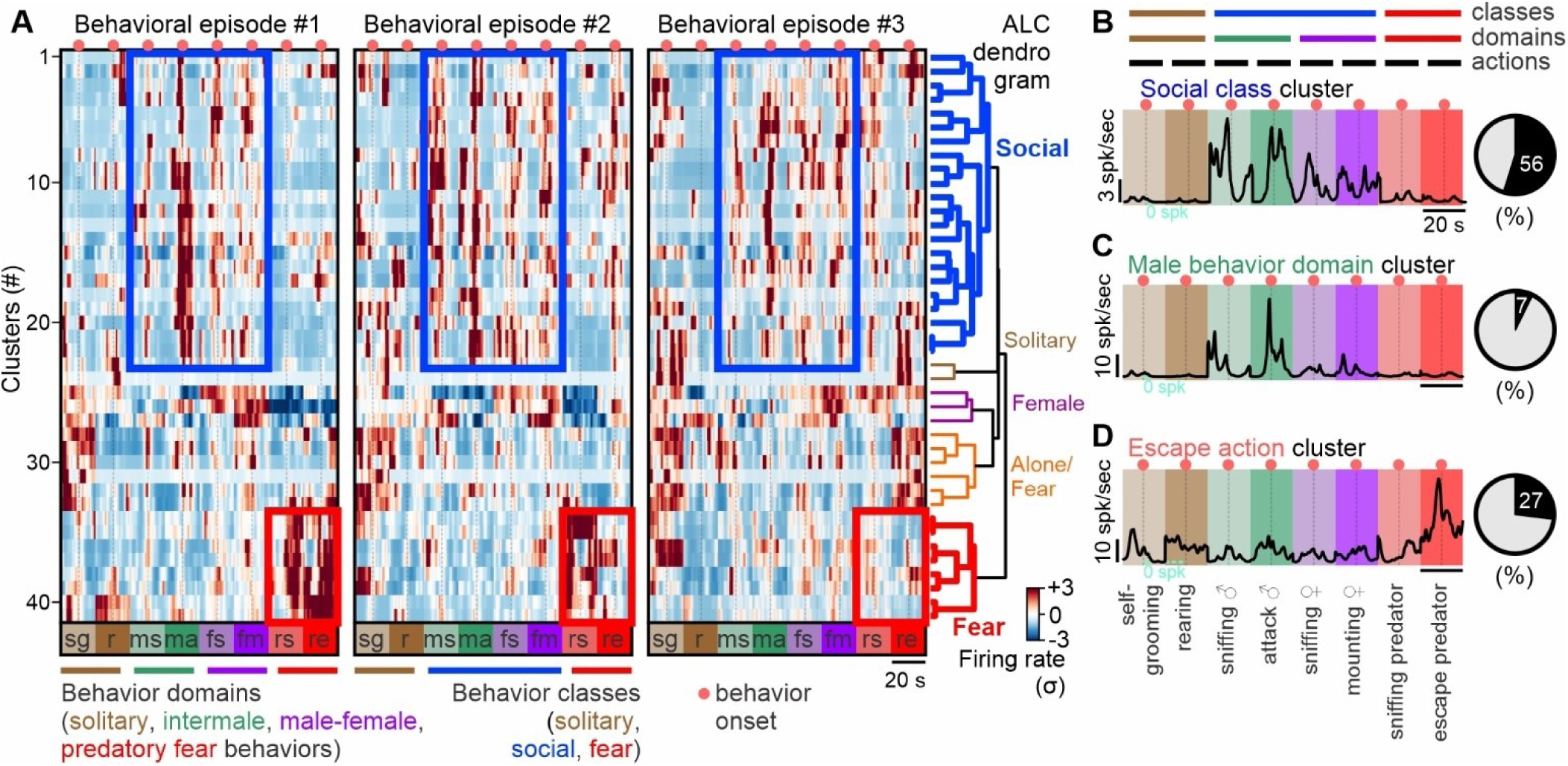
Distinct hypothalamic neuron ensembles encode social and fear behavior classes. (A) Mean activity of *k*-means clusters explaining 70% of the variance of the single unit data (Extended Data Figure 8). Clusters are sorted using agglomerative linkage clustering to generate a dendrogram whose branches have been assigned semantic labels, based on cluster activity being higher than two standard deviations in specific behavioral classes. (B) Left: Mean activity of a *k*-means cluster selected from the forty-one clusters identified and shown in (Figure 4A), exhibiting mixed selectivity, i.e., activity in diverse behavior domains – in this case, in male-male and male-female behaviors. Right: Portion of clusters showing mixed-selectivity / multi-domain activity. (C) Left: Mean activity of a *k*-means cluster selected from the forty-one clusters identified (Figure 4A), showing mixed selectivity within a behavioral domain – in this case, in intermale behaviors. Right: Portion of clusters showing mixed selectivity within a behavioral domain (alone / intermale / male-female / and fear behaviors). (D) Left: Mean activity of a *k*-means cluster selected from the forty-one clusters identified (Figure 4A), showing behavior-specific tuning – in this case, in escape response to a predator. Right: Portion of clusters showing behavior action-specific tuning. Abbreviations: sg: self-grooming, r: rearing, ms: male intruder anogenital sniffing, ma: intermale attack, fs: female intruder anogenital sniffing, fm: mounting female intruder, rs: sniffing predator, re: escape response to a predator. See also Figures S8, and S9.

In the solitary domain, the self-grooming (sg) and rearing (r) behavioral actions were included. Representative behavioral actions in the intermale domain were male intruder sniffing (ms), and attack (ma), while in the male-female domain female intruder sniffing (fs) and mounting (fm) were included. Lastly, in the predatory fear domain, the rat sniffing (rs) and escape response (re) behavior actions were included. The time points of three episodes per behavior were extracted from the experimental design illustrated in Figure 3Aa. Specifically, solitary behavior episodes were extracted from experiment (a), female-oriented behavior episodes were extracted from experiments (a) and (b), male-oriented behavior episodes were extracted from experiments (c) and (d), and predator-defence behavior episodes were extracted from experiment (e). Figure S4A shows the activity of single units across the eight above-mentioned behavioral actions, across three consecutive behavioral episodes. The activity of single units was sorted from high to low within each brain area.

To test whether separate groups of recorded units exhibit distinct behavior-correlated activity profiles, we performed *k*-means clustering of the single neuron activity (Figure S4A). The number of clusters, 41, was selected using the elbow method^35,36^; 41 clusters explained 70% of the single neuron activity data variance. The average activity of each of the 41 clusters during the 8 different domain-specific actions was plotted for each of the three consecutive episodes (Figure 4A). Agglomerative hierarchical linkage clustering (ALC) of these 41 clusters revealed five groups (i.e., dendrogram branches, Figure 4A, right, threshold for separating branches of the dendrogram was defined using the elbow method). This analysis suggested that the social and fear behavior classes are encoded by functionally distinct groups of hypothalamic neurons (Figure 4A, highlighted as blue [social] and red [predatory fear] clusters in the dendrogram). This finding was also qualitatively supported by performing non-negative matrix factorization (NMF) on the data shown in Figure S4A. However, with NMF, the distinction between social-vs. fear-correlated ensemble activity was less prominent (Figure S5A and S5B). The weaker segregation of behavioral classes through NMF likely reflects that NMF allows neurons to contribute to multiple factors (components).

While the *k*-means clusters associated with social or fear behavior classes were well separated, neurons exhibited multi-action activity, also known as mixed selectivity, within each class-specific cluster (Figure 4B). To quantify this finding, we classified a cluster as active during a specific action if its activity (spiking rate) was ≥ 2σ above its baseline activity. Accordingly, 56% of all clusters were identified to exhibit mixed selectivity within their respective behavior classes. Only seven percent of hypothalamic clusters exhibited mixed selectivity within a single behavioral domain (male- or female-directed behavior; Figure 4C), while 27% of clusters showed prominent activity during only a single action (Figure 4D). While the activity of any single cluster exhibited adaptation across episodes (Figure S4B) there was an overall positive correlation between episodes (Figure S4C) and a high-fidelity score (a measure of a cluster being identified as active in the same behavior(s) across episodes, Figure S4D). Thus, the representation of behavioral actions by different clusters is reproducible across episodes, but not rigidly fixed.

### Encoding of behavior is spatially distributed, but individual ensembles are anatomically localized

The foregoing analysis revealed that distinct neuron clusters encode social and fear behavior classes. We next sought to determine whether individual activity-defined ensembles are anatomically localized and whether they respect classical cytoarchitectonically defined nuclear boundaries.

We found that a) functionally similar clusters were distributed broadly across the hypothalamic rostrocaudal axis (Figure 5A), and b) that most clusters were composed of cells that originated from a few neighboring hypothalamic regions (Figure 5B). Re-sorting these clusters within each branch of the dendrogram according to their position along the rostro-caudal axis (Figure 5C), revealed that each behavior class was broadly represented along this axis (Figure 5C).

**Figure 5.**
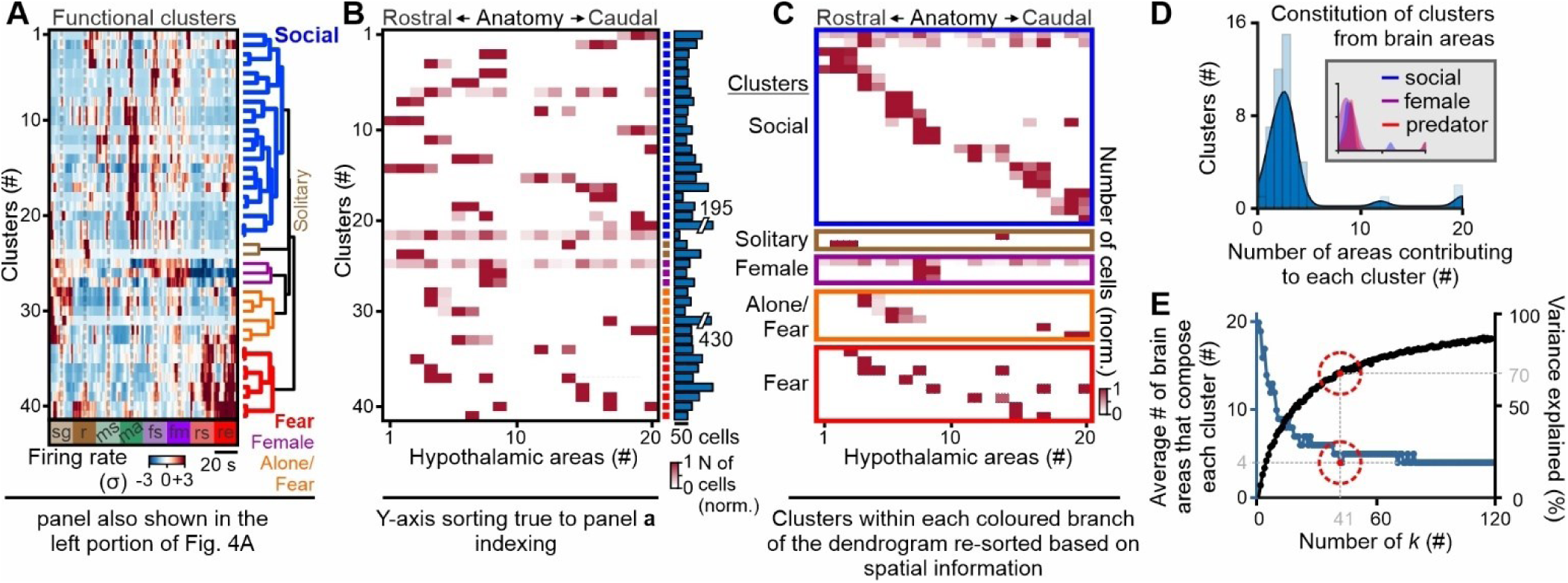
Encoding of behavior classes is spatially distributed, but single clusters are spatially localized. (A) Functional clusters of single neuron activity (panel from Figure 4A – left). (B) Spatial composition of each of the 41 functional clusters shown in panel (A). (C) Re-sorting the spatial composition of each cluster, to present clusters composed of most rostral regions, to clusters composed of most caudal regions. (D) Histogram of hypothalamic area contribution per cluster. Inset: Histogram of hypothalamic area contribution to clusters that are active in specific behaviors. (E) Spatial composition of clusters based on the choice of *k* (number of clusters selected for *k*-means clustering), reveals that the average composition of a cluster by four regions is a stable solution. Abbreviations: sg: self-grooming, r: rearing, ms: male intruder anogenital sniffing, ma: intermale attack, fs: female intruder anogenital sniffing, fm: mounting female intruder, rs: sniffing predator, re: escape response to a predator. See also Figures S9B-S9D.

The number of regions contributing to single clusters was typically four areas (Figure 5D), suggesting that most hypothalamic clusters consist of cells that originate from four neighboring hypothalamic regions. The same observation occurred when quantifying the composition of clusters that were identified as “social“, female-specific, and predatory-fear relevant (Figure 5D – inset). We investigated whether increasing the number of clusters in *k*-means leads to a reduced number of regions per cluster (Figure 5E). However, we found an asymptote at an average number of four nuclei per cluster which did not further decrease even when the number of clusters was increased up to 120 (Figure 5E). The smallest number of clusters that reached the asymptotic value of 4 regions per cluster was 41, and this model explained a majority of the variance in the data (70%). A qualitatively similar result emerged when analyzing factors identified by NMF, with closely related factors composed of neurons at different positions along the rostrocaudal axis, and heavily weighted neurons in each factor were present in neighboring hypothalamic regions (Figure S5C-S5E).

### Behavior explains a quarter of the variance of hypothalamic neuron activity

The above approaches suggest that: 1) distinct populations of neurons represent social and fear behaviors, 2) the encoding of each behavioral class is represented by a set of clusters comprised of units defined by correlated neuronal activity, which are spatially distributed across the rostro-caudal axis, and 3) individual clusters typically contained neurons from ca. four neighboring nuclei, therefore exhibiting a strong local anatomical bias in their composition. However, since most of the units in these functional clusters exhibited mixed selectivity, it remained uncertain whether specific behavior actions could be decoded from neural activity.

To determine whether individual behavioral actions could be predicted from single-unit activity, we trained decoders based on neural activity using two different machine learning algorithms (gradient boosted trees ^37^, results plotted in Figure S6A and S6B, and support vector machines ^23^, results plotted in Figure S6C-S6E). A subset of appetitive and consummatory representative behavior actions per behavioral domain were selected to address the above point. The results indicated that most, but not all, actions within each of the four different behavioral domains could be decoded from neural data significantly better than chance. In the case of male-directed attack, decoder performance was high, reaching a shuffled subtracted median F1 score of ∼0.5 (Figure S6B).

The foregoing analysis suggested that behavioral actions could be decoded from neural activity but left open the question of which features of these behaviors were encoded. To address this question, we next examined how well the firing patterns of individual units could be predicted from a weighted sum of different behavioral actions and features (features included animal’s velocity, acceleration, distance, and angle between animals). These features were computed based on pose estimation using DeepLabCut ^38^. We trained generalized linear models (GLMs) ^39,40^ to predict the activity of each neuron based on a weighted sum of selected predictor variables (design of GLMs shown in Figure 6A). This approach can: 1) identify the overall variance explained by specific predictors (Figure 6B), 2) reveal the influence of all annotated predictors on single neuron activity, by inspection of their coefficients/weights (Figure 6C), and 3) indicate whether neurons that belong to the same cluster (based on the similarity of their behavior weights – Figure 6C, see dendrogram colored branches) show any spatial bias (Figure 6D and 6E).

**Figure 6.**
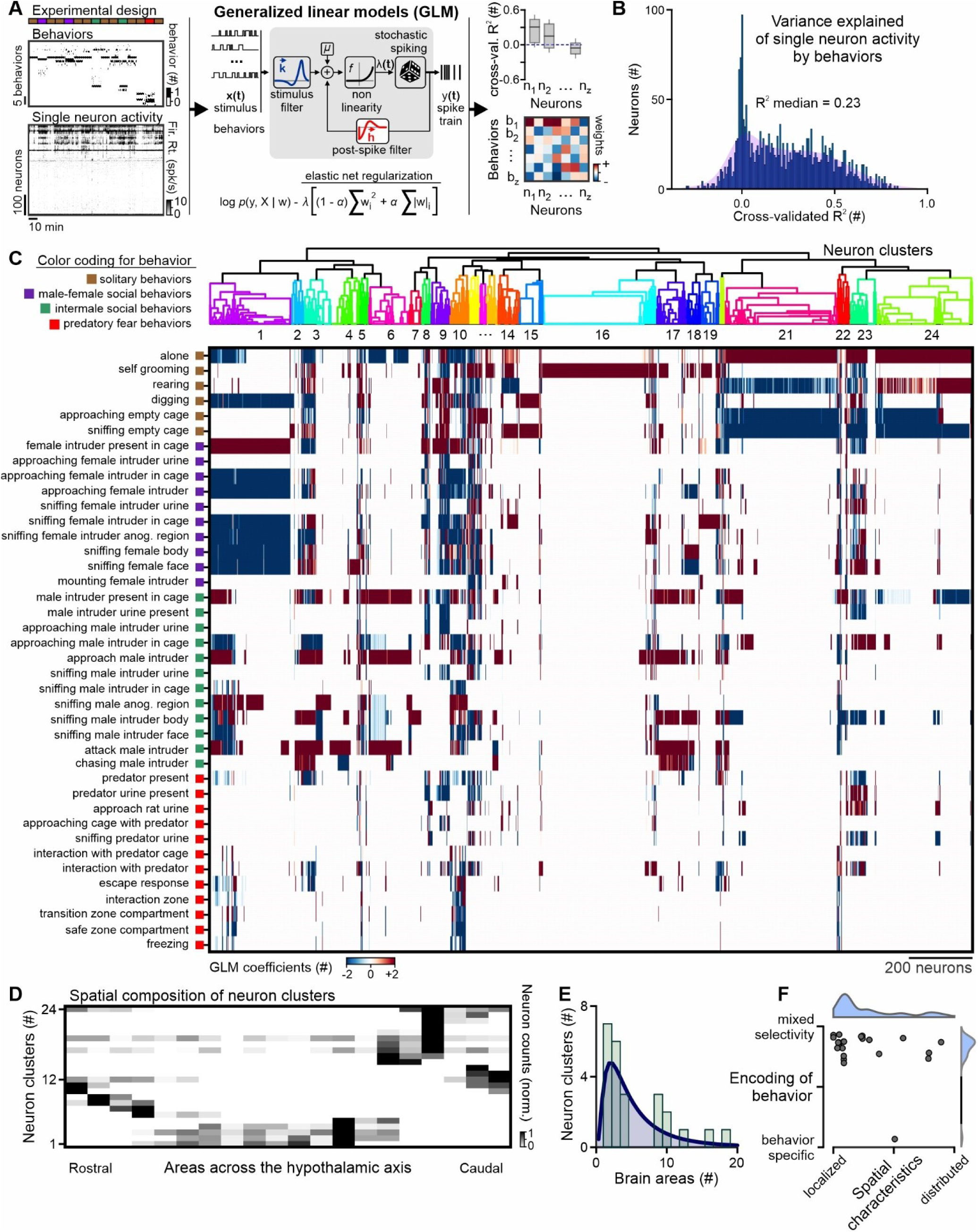
Solitary, social, and fear behaviors explain a quarter of hypothalamic neuron activity variance and clustering of encoding model weights suggest anatomical organization of functional neuron clusters. (A) Analysis design using generalized linear models (GLMs) to study the encoding of behavior by non-genetically defined mesoscale hypothalamic populations. (B) Variance explained (cross-validated, shuffled data R^2^ subtracted, R^2^) of single-neuron activity (n=6 mice, 2347 neurons) based on forty-two instinctual behaviors. (C) Agglomerative linkage clustering of behavior GLM-generated coefficients into twenty-four neural dendrogram branches/clusters. (D) Average neuron counts per brain area across the hypothalamic rostro-caudal axis across all neural branches, reveals a spatial bias of neural branch composition. (E) Histogram of hypothalamic area contribution per cluster. (F) Spatial vs. functional characteristics of neuron clusters. In box-and-whisker plots, center lines indicate medians, box edges represent the interquartile range, and whiskers extend to the minimal and maximal values. See also Figures S11, S12, and S13.

First, we sought to identify the percentage of variance explained by the four high-level behavior classes. Only 8% of the variance of single-neuron activity was accounted for by the four behavioral domains (Figure S6Fa). Next, we examined the percent of variance explained by forty behavioral actions, annotated across the four behavioral domains of interest. Behavioral actions alone explained 23% of the variance of hypothalamic neuron activity (R^2^ median = 0.23, Figure 6B). Training GLM models with predictor variables being the behavioral actions, plus motion signals (the animals’ velocity and acceleration), and proxies of sensory cues (distance and angle between animals), did not lead to an increase in the variance explained (Figure S6Ga and S6Gb).

Many different behaviors had positive and negative weights on the activity pattern of each single unit (Figure 6C), highlighting mixed selectivity as a prime mode of the hypothalamic encoding of behavior, and providing further support to the results of the *k*-means (Figure 4B) and NMF analysis (Figure S5A). Application of agglomerative clustering (ALC) to the matrix of GLM-derived behavior weights yielded 24 clusters of neurons (the elbow method was used to select the number of clusters). Each cluster (Figure 6C) contained a unique combination of neurons whose activity was influenced by a specific set of behavior weights (Figure 6C), with a near-uniform contribution of neurons from each of the six recorded mice (Figure S6H). Sorting these neurons based on their relative spatial location, however, did not generate clusters in a pattern similar to that identified by ALC, suggesting that the neurons comprising the clusters are not restricted to single hypothalamic nuclei (Figure S6I). To investigate whether the cells comprising the ALC-generated neuron clusters exhibit any restriction in their anatomic distribution, we measured the average contribution of each nucleus to each cluster (Figure 6D, rows). This revealed that clusters indeed exhibited a spatial bias, with >80% of the cells comprising each cluster originating from ∼four nearby regions (Figure 6D and 6E) – a finding consistent with the *k*-means-revealed cluster anatomical bias (Figure 5B). Also similar to the *k*-means analysis, we found that GLM-based clusters exhibited spatial bias and mixed selectivity (Figure 6F), quantified using a spatial localization index (single nucleus restricted vs. broadly distributed) and encoding entropy respectively (the degree of behavior specific vs. mixed selectivity coding; see STAR Methods section: GLM output – functional vs. spatial distribution of clusters).

## Discussion

The hypothalamus is traditionally viewed as a “low-dimensional” system in which innate social, defensive, and feeding behaviors are controlled by bulk neural activity flowing through specific pathways connecting functionally specialized anatomical subdivisions and cell types^41–43^. In contrast, the cortico-hippocampal system is thought to operate using high-dimensional, dynamic neural codes involving complex spatiotemporal patterns of population activity^44–47^. Here, we addressed whether this difference is truly biological or reflects the fact that most recording studies in the hypothalamus have used low-dimensional recording methods (e.g., fiber photometry) applied to anatomically restricted regions pre-selected based on perturbational studies. We report here what is, to our knowledge, the first unbiased high-dimensional single-unit recording of neural activity along the rostrocaudal axis of the medial hypothalamus during social and predator-defense behaviors. Our results suggest that behavioral coding in the hypothalamus is more high-dimensional, spatially distributed, dynamic, and variable than previously appreciated.

Several unexpected findings have emerged from this study. First, we observe that the initiation of these innate behaviors is (except for freezing) preceded by a hypothalamus-wide, synchronous burst of peri-onset activity, which we call an “ignition signal.” Second, we find that functionally distinct subpopulations of neurons (defined by unbiased *k*-means clustering) encode social (mating and fighting) vs. predator defense behavioral classes. Specific behavioral actions or male-vs. female-directed social behaviors are less well-discriminated by such clusters. While these clusters are reproducibly activated across multiple episodes of each behavioral class, inter-episodic variability exists in the recruitment and activity level of the neurons comprising each cluster. Nevertheless, despite this variability, many behavioral actions could be decoded from single-unit activity with high accuracy. Third, we find that the subpopulations representing social and defense behaviors are broadly distributed across the rostrocaudal length of the hypothalamus rather than spatially localized to a single nucleus or region. Fourth, we find that cluster membership draws from neurons distributed in neighboring nuclei, typically four. Nevertheless, behavior cannot be decoded from bulk neural activity based on anatomical location alone. Finally, we find that a substantial fraction (median 23%) of the single-neuron variance in activity can be explained by behavioral actions and, to a lesser degree, by behavioral domains. We consider the median R^2^ of 0.23 as a high value, as other studies which used brain-wide recordings during stereotyped behavioral tasks in head-fixed mice reported a cross-validated variance explained in single neuron activity in the range of 5-15%^48,49^.

The present study paints an overall picture that challenges prevailing views of behavioral encoding by the hypothalamus. We note that if these data had provided the very first, foundational view of hypothalamic behavioral encoding, they would not have directed subsequent functional perturbations to specific single nuclei such as VMH^11,13,22,23,50–55^ or MPOA^14,56–60^. Instead, they would have prompted the study of behavior encoding units across several neighboring nuclei. They would have shifted the focus from seeking the identification of behavior-specific cell types to understanding the properties conferred by the prime mode of activity we report here, i.e., mixed selectivity^61–65^.

Is there any way to reconcile these two seemingly opposing views of behavioral control by the hypothalamus? There are several possible explanations. One possibility is that prior, low-dimensional, anatomically restricted approaches have missed what is suggested by our silicon probe recordings: i.e., that the “units” of hypothalamic computations are anatomically distributed, and that mixed-selectivity is the prime mode of hypothalamic behavioral encoding. Recent studies suggest that even in the absence of behavior action-tuned cells, behavior can still be decoded with high accuracy, with mixed selectivity providing a basis for the encoding of numerous and diverse features in a computationally efficient manner ^61–64^. For example, nonlinear mixed selectivity enables the integration of diverse lines of information in single neurons that can elicit behavior actions based on a threshold mechanism.

A second possibility is that our silicon probe recordings have missed anatomically restricted behavior-specific subsets. This could be due to sparse sampling (we recorded on average only ∼20 units per nucleus per animal) and/or to variability in the activity of individual units across episodes and animals during a given behavior, perhaps reflecting representational drift^66,67^ or stochastic recruitment of individual units^68^. This variability may explain, in part, why a recent multi-site coarse recording of bulk calcium activity from Esr1^+^ neurons across the hypothalamus yielded reproducible, spatially distinct patterns of activity during male-vs. female-directed social behaviors, in specific nuclei previously identified as causally relevant to these behaviors by perturbation studies^69^. Alternatively, Esr1^+^ neurons may represent a “privileged,” more behavior-specific subpopulation than the “anonymous” units sampled here. A third possibility (not mutually exclusive with the second) is that the intrinsic variability in the performance or sequence of specific behavioral actions during a given activity (e.g., a male-male social interaction) may confound efforts to correlate a high-granularity annotation of these actions (43) with high-dimensional neural data (∼2400 units from 6 animals). Finally, we note that causation and representation are not the same thing: not all neural representations during a given behavior necessarily reflect a causal role for those neurons; for example, some activated units may represent efference copies of motor signals, or GABAergic neurons that serve to inhibit competing behaviors.

Further work is required to distinguish these possibilities and to arrive at an understanding of behavioral coding and function that reconciles and integrates the two views of hypothalamic computations provided by recording and perturbation strategies, respectively. At the very least, our data indicate that the application to the hypothalamus of unbiased, large-scale, anonymous single-unit recordings reveals a surprisingly complex pattern of spatiotemporal activity that does not easily map to specific nuclei or single behavior actions. It suggests that the prevailing view of the hypothalamus as a low-dimensional control system needs to be revised, or alternatively that the signals that control behavior are, to a large extent, “hidden” in the population code.

## Acknowledgments

Members of the Anderson laboratory are thanked for their advice and discussion during the preparation of this manuscript. Light-sheet imaging was performed in the Biological Imaging Facility, with the support of the Caltech Beckman Institute and the Arnold and Mabel Beckman Foundation. Special thanks to the Director of the Biological Imaging Facility at Caltech, Dr. Andres Collazo, for providing crucial experimental support and advice. We also thank Jung-Sook Chang and Xiaolin Da for experimental support, C. Chiu for laboratory management, G. Mancuso, L. Chavarria and D. R. Navarrete for administrative assistance. S.S. received support from the Wenner-Gren Foundations and by NIH K99MH131754. D.J.A. is an investigator of the Howard Hughes Medical Institute. This work was supported in part by NIH grants NS123916, MH1223612, and MH070053 to D.J.A. The content is solely the responsibility of the authors and does not necessarily represent the official views of the National Institutes of Health. All authors support an inclusive, diverse, and equitable conduct of research.

## Author contributions

S.S. conceived the study, designed and performed all experiments, wrote hardware-operating and analysis code, analyzed data, and generated the first draft of the manuscript. G.S. contributed to brain clearing, light-sheet microscopy, and silicon probe registration to the Allen Common Coordinate Framework. M.M. performed the decoding analysis using gradient boosted trees. P.P. designed decoding approaches. M.F. wrote and provided code for image registration for light-sheet microscopy data. J.K. contributed to the analysis. M.P. supervised S.S.. D.J.A. supervised S.S., contributed to the study design and analysis, and wrote the manuscript. All authors reviewed the manuscript.

## Supplemental Figures

**Figure S1.**
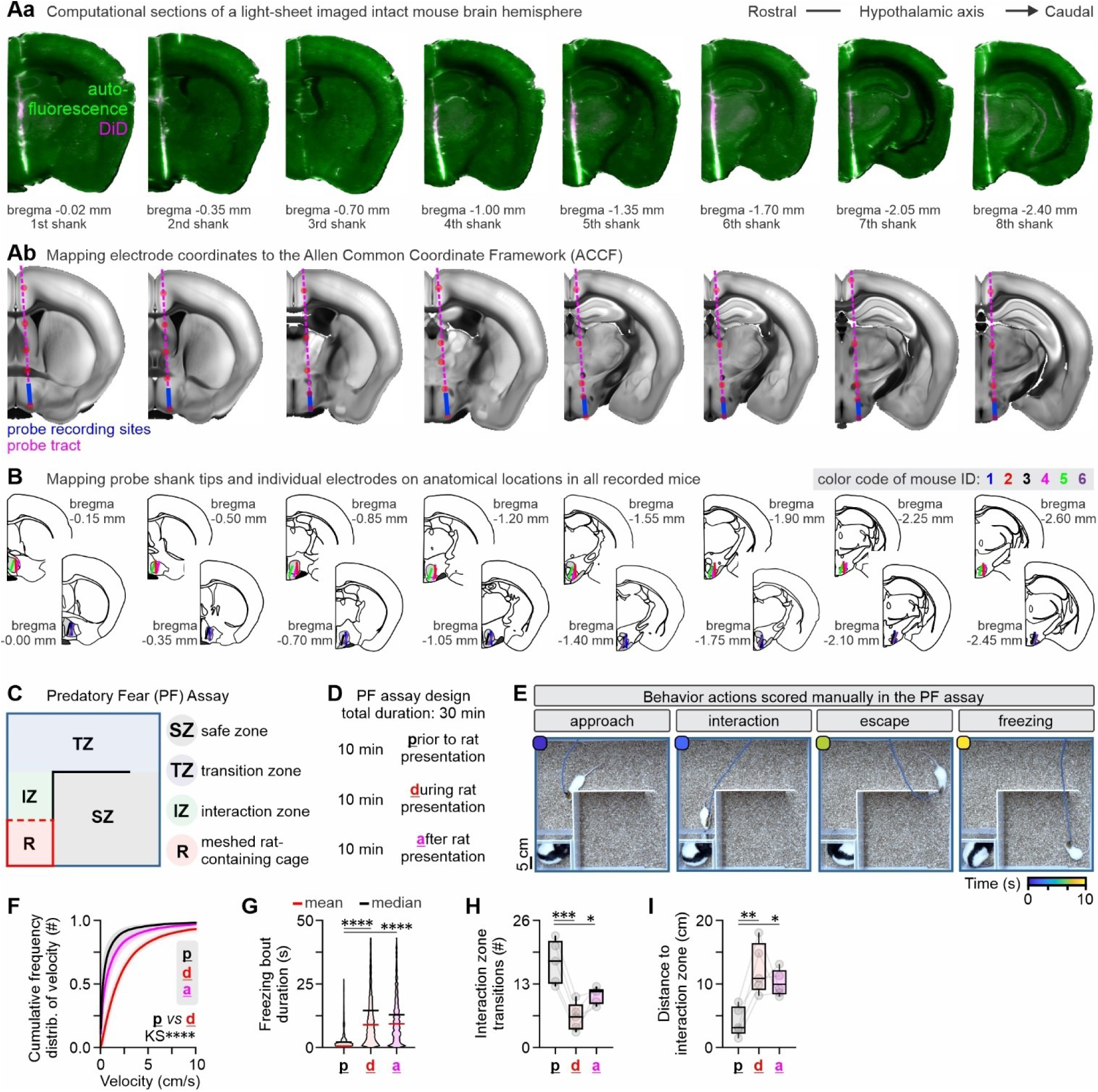
Mapping of recording sites across the rostrocaudal axis of the hypothalamus, and quantification of behavior in the predatory fear assay, related to Figures 1 and 2. (Aa) Computationally sectioned coronal views of the mouse brain that include each of the eight shanks of the chronic silicon probe implant in an example mouse brain. (Ab) Mapping the shank and the electrode-containing area of the shank to the Allen Common Coordinate Framework (example sections/mouse). (B) Mapping the overlap of all recording electrodes of each of the eight shanks across all probes/mice to the hypothalamic areas (n=6 mice). (C) Experimental design of a custom predatory fear assay, with four compartments. R: compartment containing the predator (rat), with a single, clear, and perforated side facing the compartment IZ, i.e., interaction compartment. TZ: is the transition compartment that links compartments R and SZ, the latter furthest away from the predator. (D) Timeline of the behavioral experiments with the fear condition flanked by two periods where the implanted mouse is alone in the arena. (E) Snapshots illustrating four manually annotated behaviors, specifically; approaching the predator, interacting with the predator, escape response to a predator (a high-velocity activity bout towards the safe zone compartment), and freezing behavior (observed in the freezing compartment). (F) Frequency distribution of velocity during the three conditions shown in panel (B). (G) Freezing bout duration in the three conditions presented in in panel (B). (H) Number of crosses of the interaction zone across the three conditions presented in panel (B). (I) Distance to interaction zone among the three conditions presented in panel (B). \**P* < 0.05, \*\**P* < 0.01, \*\*\**P* < 0.001, \*\*\*\**P* < 0.0001. Statistical comparisons of the data illustrated in panel (F) were performed by the Kolmogorov-Smirnov test, while comparisons of the data shown in panels (G-I) were performed through One-Way ANOVA and Dunnett’s post-hoc test was used to correct for multiple comparisons. In box-and-whisker plots, center lines indicate medians, box edges represent the interquartile range, and whiskers extend to the minimal and maximal values.

**Figure S2.**
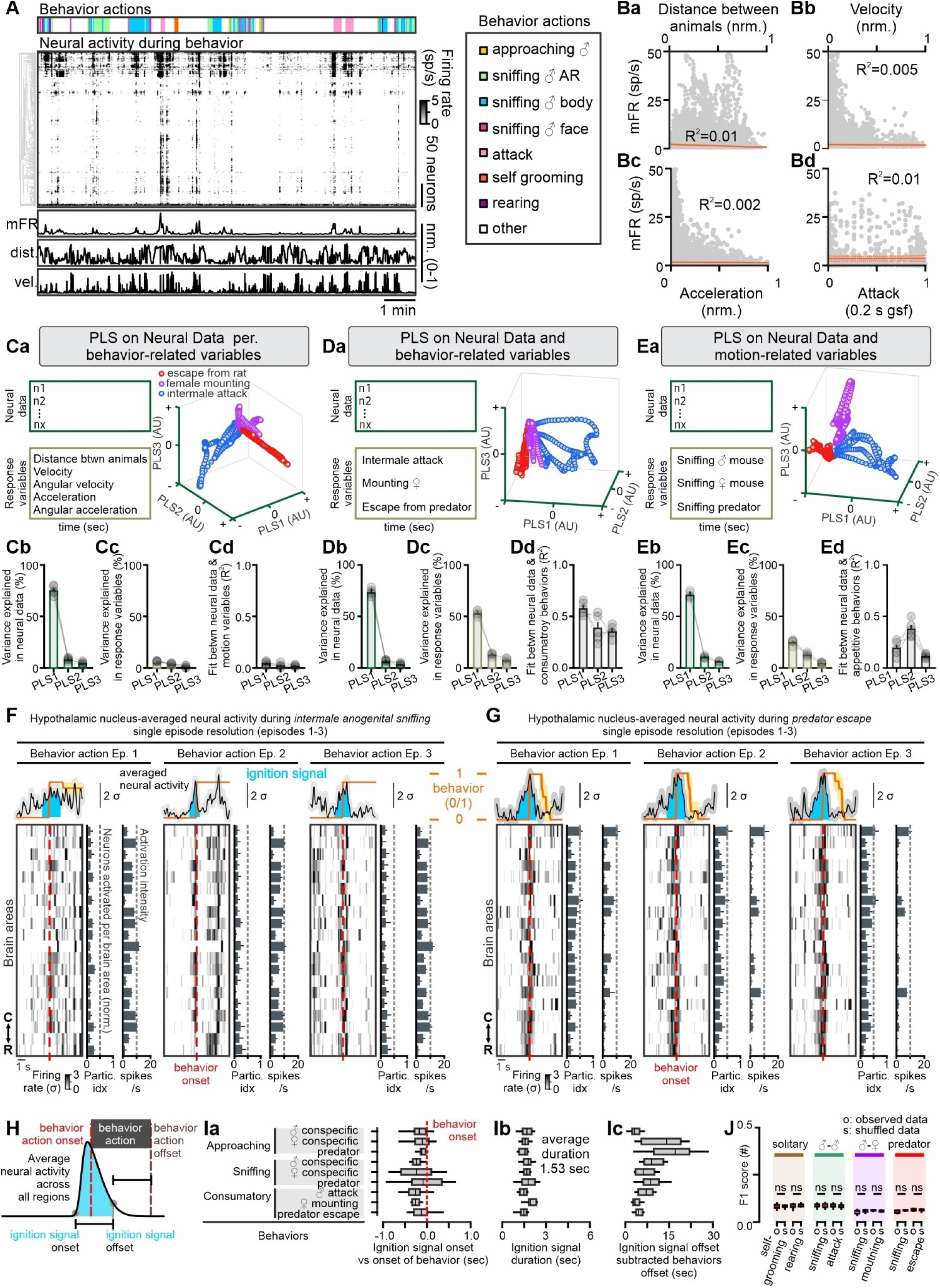
Behaviors but not motion-related variables correlate with hypothalamic neuron firing rate, and ignition signal parameter analysis, related to Figure 2. (A) Example time series data of behavior, single neuron activity, mean firing rate, distance between animals, and velocity of recorded mouse. (Ba) Linear regression between mean firing rate and distance between animals (data from six mice). (Bb) Linear regression between mean firing rate and velocity of the recorded mouse (data from six mice). (Bc) Linear regression between mean firing rate and acceleration of recorded mouse (data from six mice). (Bd) Linear regression between mean firing rate and intermale attack behavior (data from six mice). (Ca) Left: Analysis design and input to partial least-squares regression, to assess the correlation between partial least-square components between neural data and motion-related variables. Right: Neural trajectories of PLS components in the PLS space (example mouse). (Cb) Variance explained by PLS components of the single-unit neural data (n=6 mice). (Cc) Variance explained by PLS components of the motion-related variables (n=6 mice). (Cd) Fit between neural and motion-variable PLS components, indicative of a poor link between the two types of data (n=6 mice). (Da) Left: Analysis design and input to partial least-squares regression, to assess the correlation between partial least-square components between neural data and consummatory social and fear behaviors. Right: Neural trajectories of PLS components in the PLS space (example mouse). (Db) Variance explained by PLS components of the single-unit neural data(n=6 mice). (Dc) Variance explained by PLS components of the consummatory social and fear behaviors (n=6 mice). (Dd) Fit between neural and consummatory instinctual behavior-based PLS components, indicative of a link between the two types of data (n=6 mice). (Ea) Left: Analysis design and input to partial least-squares regression, to assess the correlation between partial least-square components between neural data and appetitive social and fear behaviors. Right: Neural trajectories of PLS components in the PLS space (example mouse). (Eb) Variance explained by PLS components of the single-unit neural data(n=6 mice). (Ec) Variance explained by PLS components of the appetitive social and fear behaviors (n=6 mice). (Ed) Fit between neural and appetitive social and fear behavior-based PLS components, indicative of a moderate link between the two types of data (n=6 mice). (F) Top part: Average behavior signal and mean neural activity at the peri-onset interval of sniffing a male conspecific across three behavioral episodes. Color plot part: Average neural activity per nucleus (heatmap), participation index (% of neurons activated above 2 SD from baseline activity), and average firing rate of the activated neurons (bottom, n = 6 mice, in three behavior episodes per mouse). (G) Top part: Average behavior signal and mean neural activity at the peri-onset interval of escape responses in the presence of a predator, in three behavioral episodes. Color plot part: Average neural activity per nucleus (heatmap), participation index (% of neurons activated above 2 SD from baseline activity), and average firing rate of the activated neurons (bottom, n = 6 mice, in three behavior episodes per mouse). (H) Characteristics of ignition signals include onset, offset, and duration relevant to behavior. (Ia) Quantification of the onset of ignition signals relevant to the onset of behavior (n = 6 mice, neural data averaged across three consecutive behavior instances per mouse). (Ib) Quantification of the ignition signal duration (n = 6 mice, neural data averaged across three consecutive behavior instances per mouse). (Ic) Quantification of the ignition signal offset subtracted behavior offset (n = 6 mice, neural data averaged across three consecutive behavior instances per mouse). (J) SVM decoder F1 score on behavior actions of interest based on nucleus-averaged neural activity (n = 6 mice). Abbreviations: mFR: mean firing rate all recorded neurons, dist.: distance between animals, vel.: velocity, sp/s: spikes per second, nrm.: data normalized between 0 and 1 values, 0.2 s gsf: 200 millisecond Gaussian smoothing filter applied on the spike and behavior data prior to performing linear regression. In box-and-whisker plots, center lines indicate medians, box edges represent the interquartile range, and whiskers extend to the minimal and maximal values.

**Figure S3.**
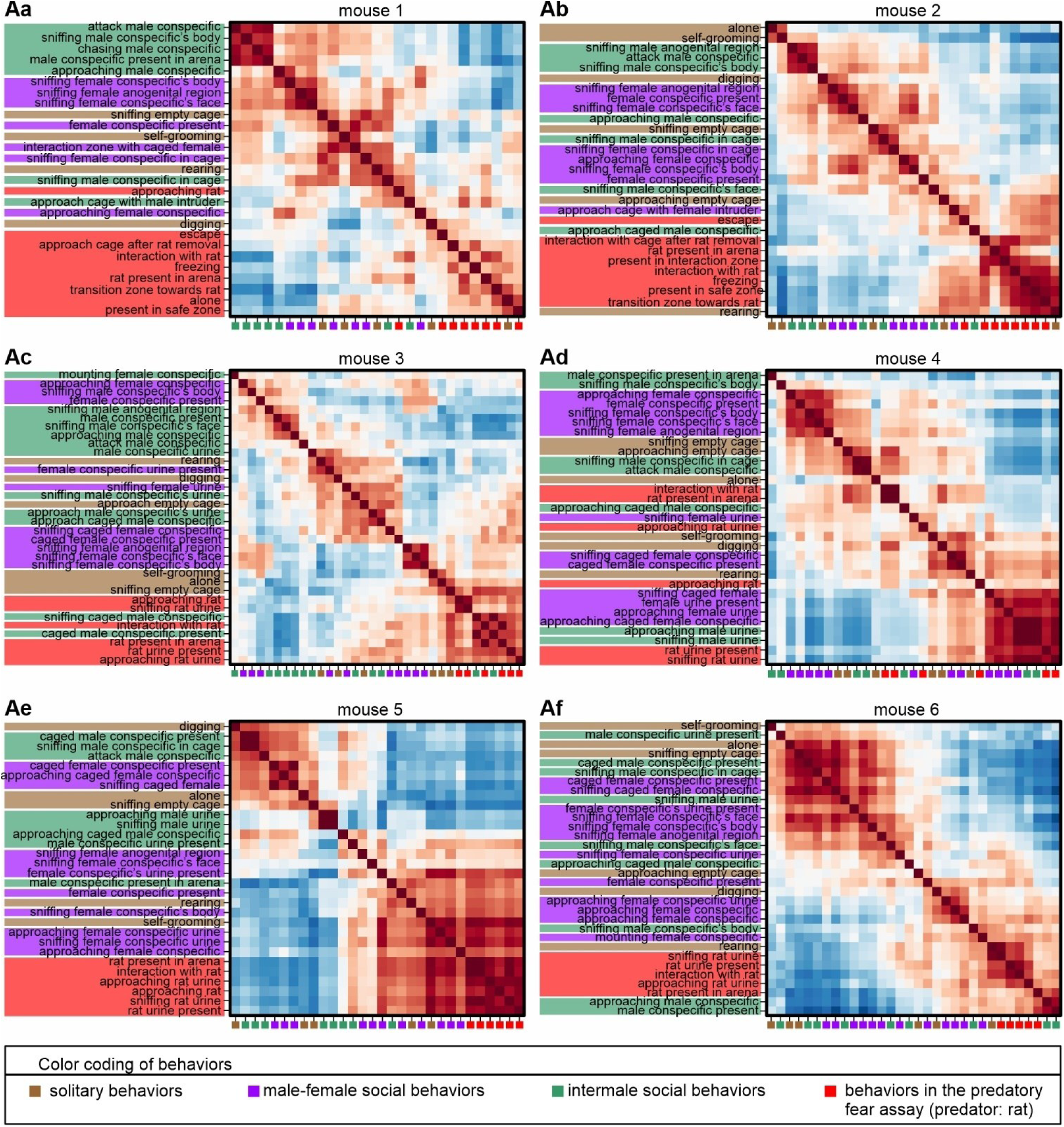
Correlation matrix of behavior for each of the recorded six mice, related to Figure 3. (Aa-Af) Correlations matrices of behavior per mouse, rows (and columns) sorted based on similarity using the rastermap algorithm. Note the segregation of fear behavior actions (marked in red) towards the end of the row labels in each matrix (with the exception of mouse #4), while other behavior are intermingled.

**Figure S4.**
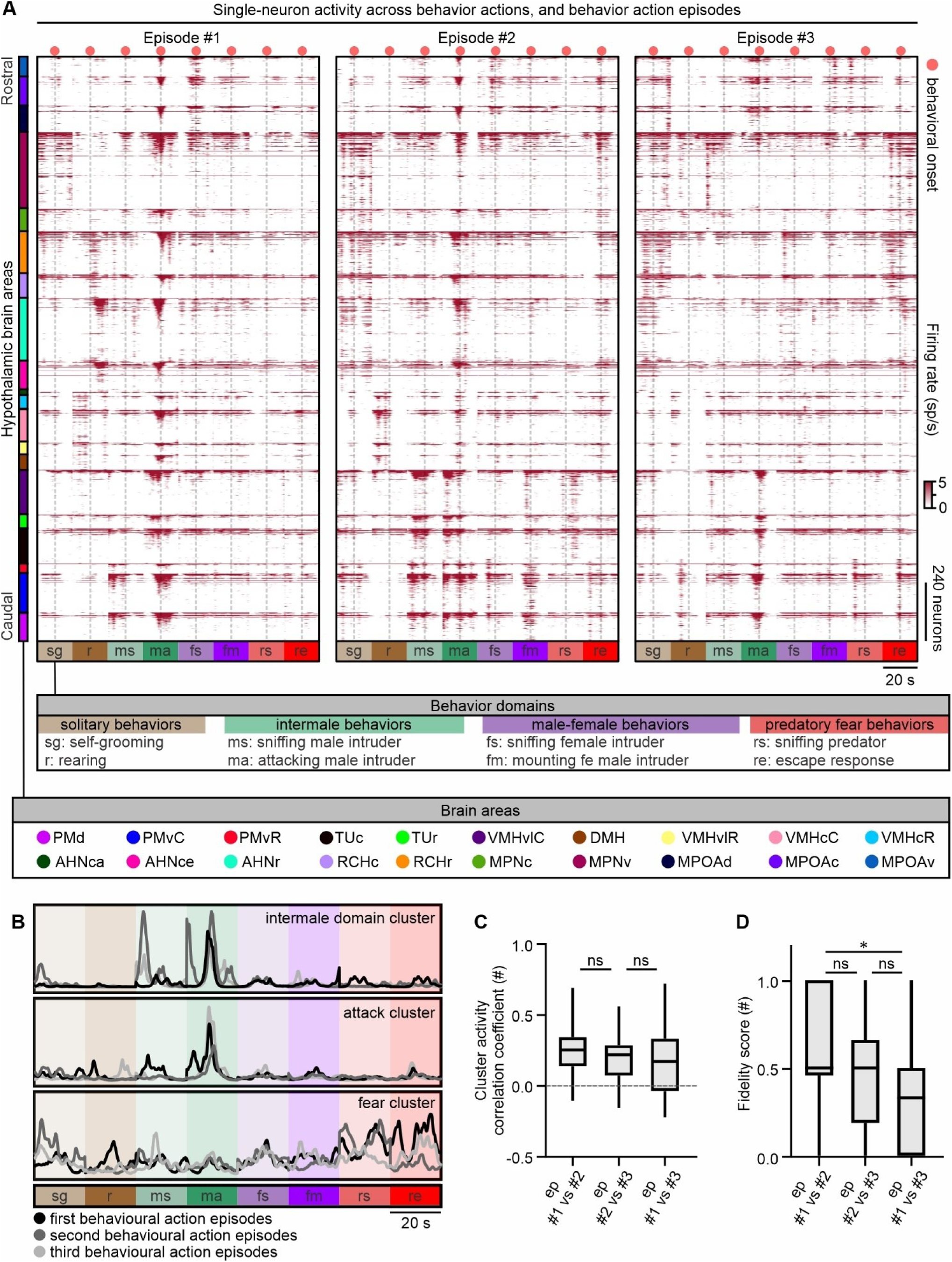
Single unit activity across four behavior domains, eight behaviors of interest, and three behavior episodes, related to Figure 4. (A) Single unit activity in eight behaviors of interest during three consecutive behavioral episodes/instances (n=6 mice, 2347 neurons). (B) Selected clusters from panel A of Figure 4, indicating the activity of each cluster across the first, second and third behavioral action episodes. (C) Correlation coefficient of average cluster activity across behavioral episodes (relevant to Figure 4A). (D) Fidelity score of cluster identity across behavior episodes; a measure that quantifies the degree to which a cluster is active to the exact same behavior(s). A cluster considered active during the expression of any single behavior, scales its activity above the threshold of two standard deviations (as compared to its baseline activity). ns: statistically not significant, \**P* < 0.05. Statistical comparisons of the data shown in panels (B, C) were performed through One-Way ANOVA and Dunnett’s post-hoc test was used to correct for multiple comparisons. In box-and-whisker plots, center lines indicate medians, box edges represent the interquartile range, and whiskers extend to the minimal and maximal values.

**Figure S5.**
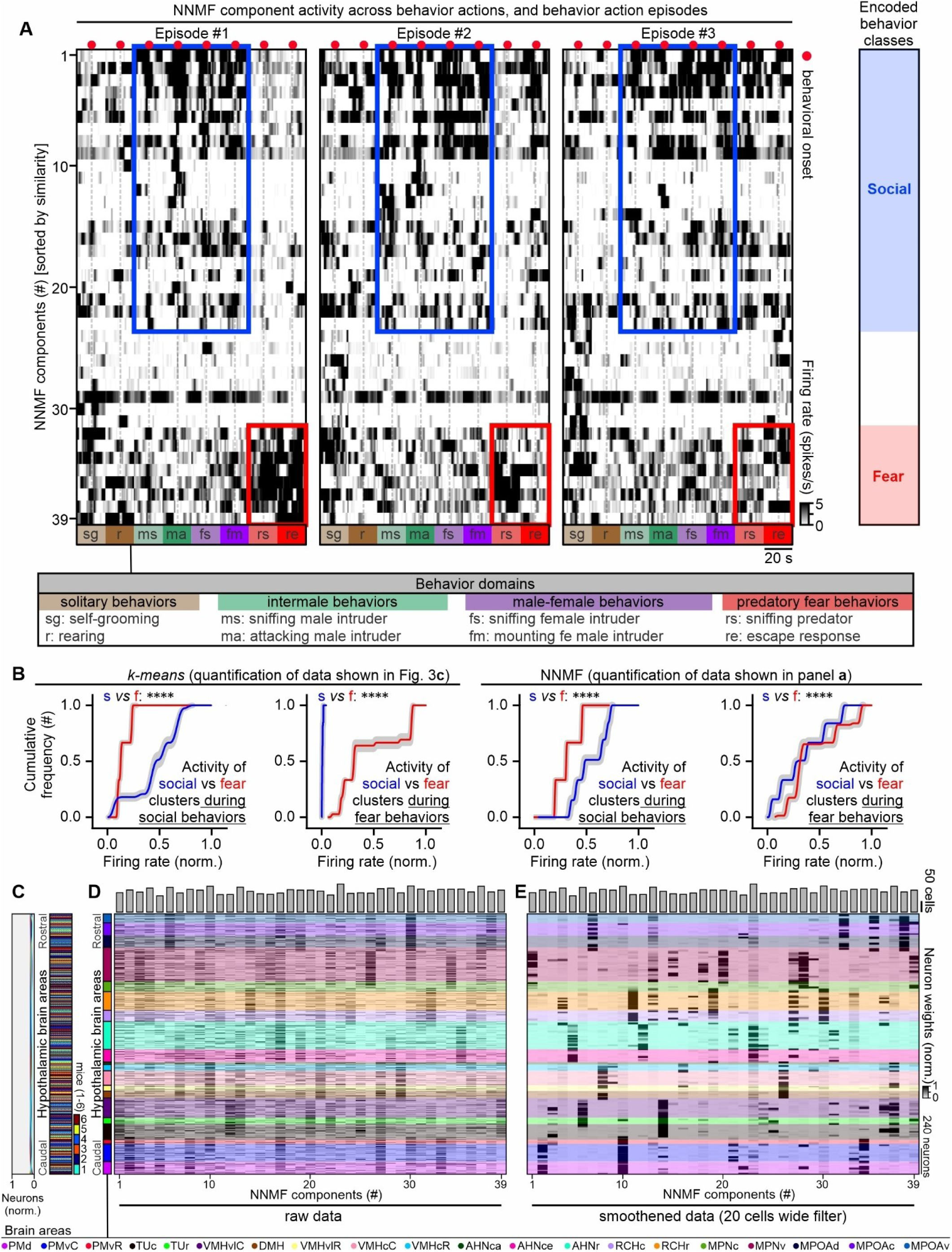
Non-negative matrix factorization (NMF) identifies ensembles with behavior class specificity and spatial bias, related to Figures 4 and 5. (A) NMF matrix factorization of the single neuron activity data shown in Figure S8, generates 39 NMF components (collectively explaining 79% of the variance of the single unit data, a number selected using the elbow method). Activity of components occurs somewhat similarly to the *k*-means clusters identified and shown in Figure 3, during the social and fear behavior classes. (B) Cumulative frequency of recorded firing rates in “social” or “fear” clusters identified in the *k*-means (Figure 4) and NMF (present figure) analysis respectively. Note that cluster activity is significantly different between conditions, but magnitude of effect is smaller in the NMF analysis. (C) Histogram (left) and raster plot (right) showing neuron origin from the six recorded mice. (D) Neuron weights of the NMF components (raw data, n=6 mice). (E) Neuron weights of the NMF components (smoothened data [Gaussian smoothing filter with twenty cells size], n=6 mice). \*\*\*\**P* < 0.0001. Statistical comparisons were performed through the Kolmogorov-Smirnov test. Abbreviations of brain areas presented in the legend of Figure 1.

**Figure S6.**
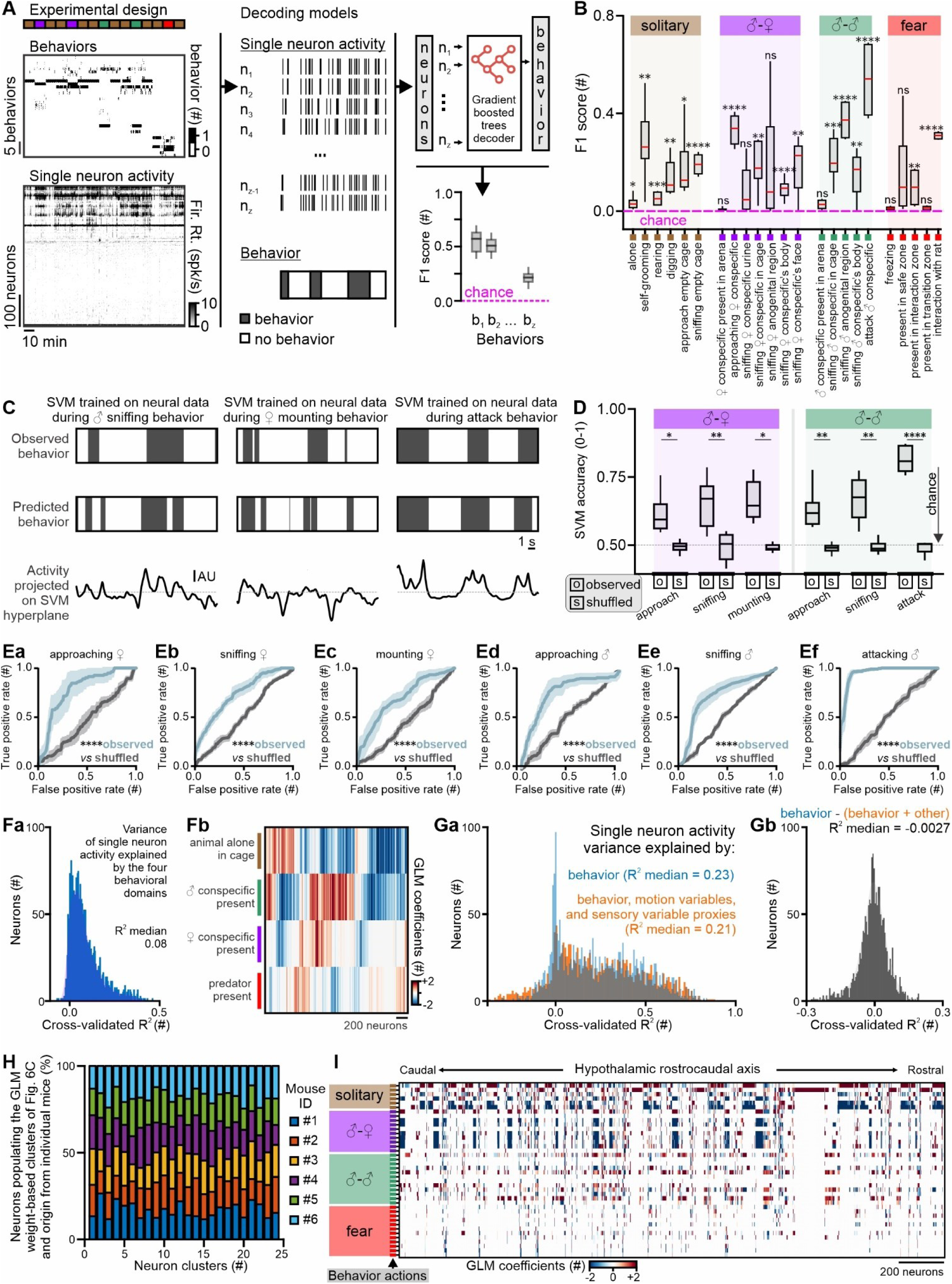
Decoding behavior from single neuron activity, and encoding models explaining single neuron activity variance based on different predictor variables,related to Figure 6. (A) Schematic of the analysis pipeline using gradient-boosted trees for binary decoding of behavior based on single unit activity. (B) Shuffled F1-score subtracted, k-fold cross-validated F1-score output of decoding behavior based on the gradient boosted trees decoders (box plots are made of points from the six recorded mice – each mouse contributing a single point unless a behavior was not expressed by an individual). In box-and-whisker plots, center lines indicate medians, box edges represent the interquartile range, and whiskers extend to the minimal and maximal values. (C) Example accuracy of three decoders on the test data of three social behaviors, and activity projected on the SVM hyperplane. (D) Decoder accuracy on social behaviors of interest (n = 6 mice). (Ea-Ef) ROC curves on the observed and shuffled data of specific social behaviors (n = 6 mice). (Fa) Variance explained (cross-validated, shuffled data R^2^ subtracted, R^2^) of single-neuron activity (n=6 mice, 2347 neurons) based on the four behavioral domains of interest, solitary, male intruder in arena, female intruder in arena, and predator in arena. Output of Poisson-GLM. (Fb) Behavioral domain coefficients on single neurons recorded across all mice (n=6 mice). (Ga) Overlap of two histograms of cross-validated R^2^ values generated by GLMs. Light blue: Variance explained (cross-validated, shuffled data R^2^ subtracted, R^2^) of single-neuron activity (n=6 mice, 2347 neurons) based on forty annotated behaviors. Orange: Variance explained (cross-validated, shuffled data R^2^ subtracted, R^2^) of single-neuron activity (n=6 mice, 2347 neurons) based on forty instinctual behaviors, two motion-related variables, i.e., animal’s velocity, animal’s acceleration, and two sensory proxy variables, i.e., distance and angle between animals. (Gb) Difference in the variance explained by behavior vs. behavior + motion-related and sensory proxy variables. Histogram was generated by subtraction of each neuron’s R^2^ generated via GLMs trained and tested only using annotated behaviors as predictor variables, against the neuron’s R^2^ generated via GLMs trained and tested using annotated behaviors and motion variables as predictors. The peak around R^2^=0 is suggestive that motion variables explain little if any of the variance of hypothalamic single neuron activity. (H) Neuron contribution from each of the recorded mice to the neural branches presented in Figure 6C. (I) Sorting neurons based on their spatial coordinates (as compared to the agglomerative linkage clustering-based sorting using behavior coefficients shown in Figure 6C). ns: statistically not significant, \**P* < 0.05, \*\**P* < 0.01, \*\*\**P* < 0.001, \*\*\*\**P* < 0.0001. In box-and-whisker plots, center lines indicate medians, box edges represent the interquartile range, and whiskers extend to the minimal and maximal values.

## STAR★METHODS

### Experimental animals

All experiments involving the use of living animals or their tissues adhered strictly to the guidelines set by the National Institutes of Health (NIH). They were conducted under the approval of both the Institutional Animal Care and Use Committee and the Institutional Biosafety Committee at the California Institute of Technology (Caltech). Swiss Webster (CFW) mice (Crl: CFW [SW]), C56BL/6NCrl mice, and Long-Evans (Crl:LE) rats were purchased from Charles River US. In all behavioral and resident-intruder assays, wild-type mice from the C56BL/6NCrl strain were used (sourced from Charles River Laboratories). Experimental animals were provided with additional enrichment and running wheels in their cages, were maintained in good health, and were monitored by the local veterinarian. Housing conditions for the mice were carefully monitored. Ventilated micro-isolator cages were maintained in an environment with stable temperature (23°C) and humidity (55-60%). A reverse light-dark cycle of 12 hours each was used, and animals were provided unlimited access to food and water. The cages were cleaned and changed once a week on a specific day, and no experiments were conducted within two days of a cage change.

### Selection of animals for experiments

*In vivo* electrophysiology data were collected from six Swiss-Webster mice (out of 11 implanted animals generated in a single cohort) and are reported here. Mice were selected for combined electrophysiology and behavior experiments based on the following criteria: (1) recovered within two days to pre-implantation baseline behaviors, (2) expressed the majority of instinctual behaviors on the experimental days, and (3) identified single neurons did not show drift across experimental sessions.

### Surgical methods

All surgical and behavioral procedures adhered to the standards set by the National Institutes of Health and received approval from the Institutional Animal Care and Use Committee at Caltech. Male Swiss Webster mice, aged between 3-4 months, were used for neural recordings during behavior. Male mice were used to avoid potential confounding effects of the estrous cycle in female mice.

Prior to surgery, mice were provided with medicated food cups (MediGel CPF, Clear H20 74-05-5022), containing carprofen, for two days. Throughout the surgical procedure, mice were kept anesthetized with isoflurane. The initial steps included removing the skin and clearing any underlying connective tissue. The tissue surrounding the skull edges was securely adhered with Vetbond (Santa Cruz Biotechnology, cat. no. sc-361931), and the skull surface was prepared by being scraped, dried, and textured.

A custom boomerang-shaped titanium head bar was affixed to the occipital bone of the skull, and a hole for the ground screw was carefully drilled, placing the screw above the right prefrontal cortex. Optibond Solo Plus (Kerr, cat No. 31514) was applied to the skull and cured under UV light. Charisma (Net32, cat. No. 66000085) was employed to establish a base for the implant and to reinforce the area around the ground screw.

With the help of a dental drill, eight 100 μm holes were made above the somatosensory cortex. These holes were designed to accommodate the eight shanks of the silicon probe, aimed at penetrating the deep brain. The entire silicon probe assembly, including the probe and headstage, was placed in the stereotax. Post-application of DiD’ (ThermoFisher Scientific, cat. no. D7757) on the posterior to the electrode surface of each shank, the probe was gently lowered into the brain at a speed of 6 μm/sec resulting in an insertion time of fifteen minutes. Higher insertion speeds negatively impacted post-surgery recovery and neuron yield per mouse.

The precise coordinates for probe insertion and the angle of attack were adapted for each mouse accommodating idiosyncratic differences in cortical vasculature identified post-drilling, and enabling the recording of the same deep brain structures. The ground wire was soldered to the ground screw and Metabond cement was used to secure the probe and headstage to the skull. The entire assembly was wrapped in Kapton Tape (ULINE S-7595), followed by a recovery period typically extending to two weeks. Post-surgery, mice were individually housed, and additional enrichment elements were provided. Daily monitoring was performed to evaluate locomotion and the expression of behaviors (self-grooming, rearing and digging). Between the second and third weeks, mice were acclimatized in test arenas to examine their natural movement and the manifestation of thigmotaxis in the open field test, among other behaviors.

### Behavioral tests

All behavioral tests were performed during the dark phase and under dim red light between 2 h after the initiation of the dark phase and three hours before the initiation of the light phase. Mice were acclimated to the testing behavior room for a minimum of one hour before testing with *ad libitum* access to food and water. Videos were recorded using the GS3-U3-51S5C-C 2/3” FLIR Grasshopper®3 USB 3.0 Color Camera, at thirty frames per second. Video acquisition was triggered through a TTL signal provided to the *trigger-*in GPIO port of the camera, by the neural data acquisition software Allego – NeuroNexus’ proprietary software for data acquisition. Timestamps for frames were collected and recorded as an auxiliary signal through the software Allego and NeuroNexus’ data acquisition box (SmartBox Pro).

### Resident–intruder (RI) test

The resident’s cage was not cleaned for a minimum of 2 days before the behavioral test. A single male or female intruder was introduced per testing session. A ten-minute habituation period was allowed and recorded, and was followed by the introduction of a freely moving intruder for ten minutes or an intruder within a perforated cage. Video and neural data acquisition continued for ten minutes following the removal of the intruder. Intruders were of the C57BL/6N background, and were either intact males, or intact females in the estrus phase of the estrous cycle (stage of cycle was defined by inspection of vaginal smears, and the cycle of individual female intruders was followed for three consecutive days prior to the day of the experiment). Behaviors during the RI test were video recorded using a top view camera, and were manually annotated using BENTO^70^ (https://github.com/neuroethology/bentoMAT).

### Predatory fear (PF) behavioral assay

A custom four-chamber arena was used to present a caged-predator. All but the front sides of the cage of the predator were opaque white colored, with the exception of the front side that was clear and the wall was perforated evenly with square holes five square millimetres in size placed fifteen millimetres apart (center to center). This design aims to allow diffusion of pheromonal cues that may affect the behavior of pray animals. Other than the chamber containing the predator, the arena includes three other chambers: i) the interaction zone chamber that allows the mouse to sniff the predator and interact with the cage before, during and after the presence of the rat in the arena. ii) The transition zone chamber that allows the mouse to move between the proximal and furthest away chambers from the rat, and iii) the safe zone chamber, that places a large distance between the mouse and the predator. The validation parameters of this custom experimental design are presented in extended data figure four.

### Behavior annotations and quantification of behavior

Behavior videos were manually annotated using BENTO^70^ (freely available software package: https://github.com/neuroethology/bentoMAT). A total of forty-three behaviors during the resident intruder and predatory fear assays were annotated. The behavioral videos were loaded into MATLAB, and a trained individual blind to the experimental design annotated the videos frame-by-frame for all behavior actions. Behavior annotations were then further edited by an expert (author: S.S.) who was not blind to the experiment conditions. Imaging frames were spatially downsampled by a factor of two, a modification that we recognize to not influence the identification of the behaviors of interest and that facilitates smoother processing of the videos by BENTO^70^ in MATLAB. Behavior features including velocity, acceleration, distance, and angle between animals were calculate using DeepLabCut^38^, and custom MATLAB routines.

### *In vivo* electrophysiology

*In vivo* electrophysiology recordings were performed in freely moving mice, using chronic silicon probe implants. All extracellular recordings were conducted across the rostrocaudal hypothalamic axis of the left hemisphere. Recordings were performed using a 256-channel custom-built for this specific project silicon probe, the A8×32-Poly2-7.5mm-25s-325-177 NeuroNexus probe, now commercially available. This specific probe includes eight shanks evenly spaced by 325 μm, and each shank at its lower end contains two columns of electrodes, each electrode with a 25 μm distance from the next. The recording area of each electrode was designed to be 25 μm^2^. The length of each shank was 7.5 mm to enable the probe to reach the model ventral nuclei of the deep brain in the hypothalamus.

Electrophysiology data was collected with Allego (proprietary software of NeuroNexus, recently updated to Radiens [https://www.neuronexus.com/products/software/radiens/]), at 20 kHz sampling frequency. Kilosort 2.5 was used for spike sorting ^30^, and we refined its output with manual curation of the results through phy: https://github.com/kwikteam/phy. Neurons that exhibited drift or units that had multiunit activity were excluded. To assess the neural recordings and perform basic related analysis prior to exploring the data relevant to behavior, we used MATLAB code that can be found here: https://github.com/cortex-lab/spikes. The quality of recordings was assessed through quantification of the event rate, i.e., the frequency of spikes that are higher than the background noise level. We considered an ’event’ to be any occurrence that matches in time (less than one millisecond) and space (50 millimeters in radius) and that has a strength that is at least six times greater than the median absolute deviation (MAD) ^71^. Secondly, for unit quality assessment, we computed and plotted the signal-to-noise ratio (SNR) for each event. Event detection (spikes), spike amplitudes, and firing rates were further inspected with custom MATLAB routines (https://github.com/jenniferColonell/Neuropixels_evaluation_tools).

### Whole-brain Imaging and Anatomical Registration of Probe Electrodes

Brains that had been cleared, were imaged using the LaVision BioTec UltraMicroscope II in dibenzylether at two different wavelengths: 488 nm (which creates green autofluorescence) and 632 nm (which allows for the visualization of electrode tracks through the DiD dye, a fixable, lipophilic far-red dye that remains stable even in dehydrated conditions). The imaging was done using an Olympus MVPLAPO 2X/0.5 objective, a 5 µm step size, approximately 5 µm pixel size (at 0.8X magnification), and a single light sheet focus. The imaging only included the hemisphere of the brain where the electrode tracks were located, and it was carried out in a sagittal orientation.

Images obtained from the green and far-red channels were subsequently downsampled to a 25 µm resolution by means of averaging. The green channel was then matched to the Allen Brain Atlas CCFv3 autofluorescence atlas with a 25 µm resolution using a two-step process: an initial affine transformation followed by a nonlinear transformation carried out in Elastix. Previous studies have demonstrated that this method has an average registration error of about four voxels (or 100 µm). The same transformation process was also applied to the red channel.

Electrode tracks were traced in the registered space using a custom R code that is based on AllenCCF (https://github.com/cortex-lab/allenCCF). This was done after electrophysiological recordings were made following the placement of a chronic implant that targeted the full anterior-posterior length of the hypothalamus.

For the final analysis, brain regions were organized into twenty distinct groups based on the Allen Brain Atlas hierarchy. The specific location of each unit was determined based on where that unit produced the highest amplitude on the electrode. The location along each electrode was transformed into the Allen Brain Atlas space according to the orientation and position of the traced electrode track for each recording. The final part of allocating each electrode to a hypothalamic region was performed manually, to further improve the output of the earlier-mentioned process. For each mouse, a unique spatial map of electrodes/hypothalamic brain areas was defined and was later used to assign individual extracellular units to specific anatomical regions.

### *k*-means clustering

The variance explained by a clustering solution in the context of the *k*-means algorithm reflects the proportion of the total variance in the dataset that is “captured” by the centroids of the clusters. In other words, it indicates how well the centroids represent the data. For computing the variance explained is we used the sum of squared distances between data points and their assigned cluster centroids. The more the variance explained, the closer the data points are to their respective centroids, and hence the better the clustering solution. Specifically, we calculated within-cluster sum of squares (WCSS). After performing *k*-means clustering we obtained the cell indices, which provide the cluster assignments for each data point, and the centroids for the clusters. The sum of squared distances of each point to its cluster centroid gives us the WCSS. We then iterated over all data point. For each point, we calculated the squared Euclidean distance between the point and its assigned centroid and summed these squared distances. We then computed the total sum of squares (TSS). i.e., the sum of squared distances of each point from the overall mean of the data. The variance explained by the clustering solution can be found by subtracting the WCSS from the TSS and then dividing by the TSS. Note that as the number of clusters *k* increases, the WCSS tends to decrease (each point can be closer to its centroid), but there’s usually a diminishing return. This is often visualized with an “elbow plot,” where the rate of decrease in WCSS slows down after a certain number of clusters, indicating a potential good choice for k.

### GLM output – functional vs. spatial distribution of clusters

Neuronal cluster data was analyzed to determine its relationship with four predefined behavioral categories. For each cluster, its activity was averaged across specific row ranges corresponding to each category. Negative activity was disregarded, and data was normalized using the z-score to mitigate outlier effects. The entropy was computed for each cluster based on its normalized activity distribution across behavioral categories. Entropy was utilized to quantify the distribution of each neuronal cluster’s activity across the four behavioral categories. After averaging the activity for each category and normalizing these averages to form a probability distribution, the entropy was computed using the formula for a discrete probability distribution. This measure provided insights into whether a cluster’s activity predominantly aligned with specific behavioral categories (low entropy) or was distributed across multiple categories (high entropy). For spatial analysis, a “center of mass” approach was employed to gauge the distribution of each cluster’s activity across 20 assumed brain regions. Both the functional (behavioral category-based) and spatial distributed-ness of each cluster were then visualized using a scatter plot.

### Decoding Analysis

#### Gradient boosted trees

neuronal data (spike rates, binned at 100 ms) was smoothened through convolution with a Gaussian kernel of 10 timesteps (1 second). The data to the decoder was then made recurrent by creating a temporal window of size ten at each given timestamp. The data was then flattened before decoding. The data was analyzed using five-fold cross-validation. Each behavior was decoded individually based on the activity of all neurons. Folds were created by determining the onsets of the behavior and then creating a temporal window of the length of the behavior before its onset. All events were concatenated, and cross-validation was performed on the resulting data. The decoding was performed using gradient-boosted trees ^72,73^ (depth of 3, 50 estimators, lr = 0.01). Performance was evaluated by calculating the cross validated, shuffled data subtracted F1-score per behavior. For each behavior, we calculated the maximum performance within a specific temporal jitter (10 timesteps). We then averaged performances across animals.

#### Support Vector Machines

To assess if single-neuron activity holds information about specific social actions, we trained a unique binary linear decoder for each behavioral action to predict whether or not the behavior was taking place. To ensure the behavior’s representations weren’t mixed with the representations of the intruder’s gender, we created separate decoders for behaviors directed towards males and females. When the behavior of interest was not being exhibited, we under-sampled the negative training data to balance the number of positive and negative training examples. We then used these binary labels alongside their matching activity vectors to train a linear SVM decoder.

## Code Availability

The code used for analysis in this study is available upon request from the first author.

## References

1. Lin, X.D., Wang, S.Q., Yu, X.D., Liu, Z.G., Wang, F., Li, W.T., Cheng, S.H., Dai, Q.Y., and Shi, P. (2015). High-throughput mapping of brain-wide activity in awake and drug-responsive vertebrates. Lab on a chip 15, 680–689. 10.1039/c4lc01186d.

2. Litvina, E., Adams, A., Barth, A., Bruchez, M., Carson, J., Chung, J.E., Dupre, K.B., Frank, L.M., Gates, K.M., Harris, K.M., et al. (2019). BRAIN Initiative: Cutting-Edge Tools and Resources for the Community. Journal of Neuroscience 39, 8275–8284. 10.1523/Jneurosci.1169-19.2019.

3. Perich, M.G., and Rajan, K. (2020). Rethinking brain-wide interactions through multi-region ’network of networks’ models. Current opinion in neurobiology 65, 146–151. 10.1016/j.conb.2020.11.003.

4. Edelman, B.J., and Macé, E. (2021). Functional ultrasound brain imaging: Bridging networks, neurons, and behavior. Current Opinion in Biomedical Engineering 18. 10.1016/j.cobme.2021.100286.

5. Schrodel, T., Prevedel, R., Aumayr, K., Zimmer, M., and Vaziri, A. (2013). Brain-wide 3D imaging of neuronal activity in Caenorhabditis elegans with sculpted light. Nature methods 10, 1013–1020. 10.1038/nmeth.2637.

6. Brodt, S., Gais, S., Beck, J., Erb, M., Scheffler, K., and Schönauer, M. (2018). Fast track to the neocortex: A memory engram in the posterior parietal cortex. Science 362, 1045-+. 10.1126/science.aau2528.

7. Chersi, F., and Burgess, N. (2015). The Cognitive Architecture of Spatial Navigation: Hippocampal and Striatal Contributions. Neuron 88, 64–77. 10.1016/j.neuron.2015.09.021.

8. Doeller, C.F., Barry, C., and Burgess, N. (2010). Evidence for grid cells in a human memory network. Nature 463, 657–U687. 10.1038/nature08704.

9. Grieco, S.F., Holmes, T.C., and Xu, X.M. (2023). Probing neural circuit mechanisms in Alzheimer’s disease using novel technologies. Mol Psychiatr. 10.1038/s41380-023-02018-x.

10. Tomé, D.F., Sadeh, S., and Clopath, C. (2022). Coordinated hippocampal-thalamic-cortical communication crucial for engram dynamics underneath systems consolidation. Nature communications 13. 10.1038/s41467-022-28339-z.

11. Stagkourakis, S., Spigolon, G., Liu, G., and Anderson, D.J. (2020). Experience-dependent plasticity in an innate social behavior is mediated by hypothalamic LTP. Proceedings of the National Academy of Sciences of the United States of America 117, 25789–25799. 10.1073/pnas.2011782117.

12. Stagkourakis, S., Spigolon, G., Williams, P., Protzmann, J., Fisone, G., and Broberger, C. (2018). A neural network for intermale aggression to establish social hierarchy. Nature neuroscience 21, 834-+. 10.1038/s41593-018-0153-x.

13. Lin, D., Boyle, M.P., Dollar, P., Lee, H., Lein, E.S., Perona, P., and Anderson, D.J. (2011). Functional identification of an aggression locus in the mouse hypothalamus. Nature 470, 221–226. 10.1038/nature09736.

14. Karigo, T., Kennedy, A., Yang, B., Liu, M.Y., Tai, D., Wahle, I.A., and Anderson, D.J. (2021). Distinct hypothalamic control of same- and opposite-sex mounting behaviour in mice (vol 589, pg 258, 2021). Nature 589, E9–E9. 10.1038/s41586-020-03143-1.

15. Kunwar, P.S., Zelikowsky, M., Remedios, R., Cai, H.J., Yilmaz, M., Meister, M., and Anderson, D.J. (2015). Ventromedial hypothalamic neurons control a defensive emotion state. eLife 4. 10.7554/eLife.06633.

16. Adhikari, A., Lerner, T.N., Finkelstein, J., Pak, S., Jennings, J.H., Davidson, T.J., Ferenczi, E., Gunaydin, L.A., Irzabekov, J.J.M., Ye, L., et al. (2015). Basomedial amygdala mediates top-down control of anxiety and fear. Nature 527, 179-+. 10.1038/nature15698.

17. Kim, J., Zhang, X.Y., Muralidhar, S., LeBlanc, S.A., and Tonegawa, S. (2017). Basolateral to Central Amygdala Neural Circuits for Appetitive Behaviors. Neuron 93, 1464-+. 10.1016/j.neuron.2017.02.034.

18. Scott, N., Prigge, M., Yizhar, O., and Kimchi, T. (2015). A sexually dimorphic hypothalamic circuit controls maternal care and oxytocin secretion. Nature 525, 519-+. 10.1038/nature15378.

19. Stagkourakis, S., Spigolon, G., Liu, G., and Anderson, D.J. (2020). Experience-dependent plasticity in an innate social behavior is mediated by hypothalamic LTP. Proceedings of the National Academy of Sciences of the United States of America 117, 25789–25799. 10.1073/pnas.2011782117.

20. Mahadevia, D., Saha, R., Manganaro, A., Chuhma, N., Ziolkowski-Blake, A., Morgan, A.A., Dumitriu, D., Rayport, S., and Ansorge, M.S. (2021). Dopamine promotes aggression in mice via ventral tegmental area to lateral septum projections. Nature communications 12. 10.1038/s41467-021-27092-z.

21. Falkner, A.L., Wei, D., Song, A., Watsek, L.W., Chen, I., Chen, P., Feng, J.E., and Lin, D. (2020). Hierarchical Representations of Aggression in a Hypothalamic-Midbrain Circuit. Neuron 106, 637–648 e636. 10.1016/j.neuron.2020.02.014.

22. Lee, H., Kim, D.W., Remedios, R., Anthony, T.E., Chang, A., Madisen, L., Zeng, H., and Anderson, D.J. (2014). Scalable control of mounting and attack by Esr1+ neurons in the ventromedial hypothalamus. Nature 509, 627–632. 10.1038/nature13169.

23. Remedios, R., Kennedy, A., Zelikowsky, M., Grewe, B.F., Schnitzer, M.J., and Anderson, D.J. (2017). Social behaviour shapes hypothalamic neural ensemble representations of conspecific sex. Nature 550, 388-+. 10.1038/nature23885.

24. Nair, A., Karigo, T., Yang, B., Ganguli, S., Schnitzer, M.J., Linderman, S.W., Anderson, D.J., and Kennedy, A. (2023). An approximate line attractor in the hypothalamus encodes an aggressive state. Cell 186, 178–193 e115. 10.1016/j.cell.2022.11.027.

25. Asaba, A., Nomoto, K., Osakada, T., Matsuo, T., Kobayakawa, K., Kobayakawa, R., Touhara, K., Mogi, K., and Kikusui, T. (2022). Prelimbic cortex responds to male ultrasonic vocalizations in the presence of a male pheromone in female mice. Frontiers in neural circuits 16. 10.3389/fncir.2022.956201.

26. Ishii, K.K., and Touhara, K. (2019). Neural circuits regulating sexual behaviors via the olfactory system in mice. Neuroscience research 140, 59–76. 10.1016/j.neures.2018.10.009.

27. Itakura, T., Murata, K., Miyamichi, K., Ishii, K.K., Yoshihara, Y., and Touhara, K. (2022). A single vomeronasal receptor promotes intermale aggression through dedicated hypothalamic neurons. Neuron 110, 2455–2469 e2458. 10.1016/j.neuron.2022.05.002.

28. Renier, N., Wu, Z.H., Simon, D.J., Yang, J., Ariel, P., and Tessier-Lavigne, M. (2014). iDISCO: A Simple, RapidMethod to Immunolabel Large Tissue Samples for Volume Imaging. Cell 159, 896–910. 10.1016/j.cell.2014.10.010.

29. Wang, Q.X., Ding, S.L., Li, Y., Royall, J., Feng, D., Lesnar, P., Graddis, N., Naeemi, M., Facer, B., Ho, A., et al. (2020). The Allen Mouse Brain Common Coordinate Framework: A 3D Reference Atlas. Cell 181, 936-+. 10.1016/j.cell.2020.04.007.

30. Steinmetz, N.A., Aydin, C., Lebedeva, A., Okun, M., Pachitariu, M., Bauza, M., Beau, M., Bhagat, J., Bohm, C., Broux, M., et al. (2021). Neuropixels 2.0: A miniaturized high-density probe for stable, long-term brain recordings. Science 372, 258-+. 10.1126/science.abf4588.

31. Vinck, M., Batista-Brito, R., Knoblich, U., and Cardin, J.A. (2015). Arousal and Locomotion Make Distinct Contributions to Cortical Activity Patterns and Visual Encoding. Neuron 86, 740–754. 10.1016/j.neuron.2015.03.028.

32. Dadarlat, M.C., and Stryker, M.P. (2017). Locomotion Enhances Neural Encoding of Visual Stimuli in Mouse V1. Journal of Neuroscience 37, 3764–3775. 10.1523/Jneurosci.2728-16.2017.

33. Christensen, A.J., and Pillow, J.W. (2022). Reduced neural activity but improved coding in rodent higher-order visual cortex during locomotion. Nature communications 13, 1676. 10.1038/s41467-022-29200-z.

34. Stringer C., Z.L., Syeda A., Du F., Kesa M., & Pachitariu M. (2023). Rastermap: a discovery method for neural population recordings. bioRxiv. 10.1101/2023.07.25.550571.

35. Goutte, C., Toft, P., Rostrup, E., Nielsen, F.Å., and Hansen, L.K. (1999). On clustering fMRI time series. NeuroImage 9, 298–310. DOI 10.1006/nimg.1998.0391.

36. Thorndike, R.L. (1953). Who Belongs in the Family? Psychometrika 18, 267–276.

37. Viejo, G., Cortier, T., and Peyrache, A. (2018). Brain-state invariant thalamo-cortical coordination revealed by nonlinear encoders. PLoS computational biology 14. 10.1371/journal.pcbi.1006041.

38. Mathis, A., Mamidanna, P., Cury, K.M., Abe, T., Murthy, V.N., Mathis, M.W., and Bethge, M. (2018). DeepLabCut: markerless pose estimation of user-defined body parts with deep learning. Nature neuroscience 21, 1281–1289. 10.1038/s41593-018-0209-y.

39. Weber, A.I., and Pillow, J.W. (2017). Capturing the Dynamical Repertoire of Single Neurons with Generalized Linear Models. Neural computation 29, 3260–3289. 10.1162/neco_a_01021.

40. Park, I.M., Meister, M.L.R., Huk, A.C., and Pillow, J.W. (2014). Encoding and decoding in parietal cortex during sensorimotor decision-making. Nature neuroscience 17, 1395–1403. 10.1038/nn.3800.

41. Mei, L., Osakada, T., and Lin, D. (2023). Hypothalamic control of innate social behaviors. Science 382, 399–404. 10.1126/science.adh8489.

42. Hahn, J.D., Fink, G., Kruk, M.R., and Stanley, B.G. (2019). Editorial: Current Views of Hypothalamic Contributions to the Control of Motivated Behaviors. Frontiers in systems neuroscience 13. 10.3389/fnsys.2019.00032.

43. Dulac, C., O’Connell, L.A., and Wu, Z. (2014). Neural control of maternal and paternal behaviors. Science 345, 765–770. 10.1126/science.1253291.

44. Basu, J., and Siegelbaum, S.A. (2015). The Corticohippocampal Circuit, Synaptic Plasticity, and Memory. Cold Spring Harb Perspect Biol 7. 10.1101/cshperspect.a021733.

45. Aoi, M.C., Mante, V., and Pillow, J.W. (2020). Prefrontal cortex exhibits multidimensional dynamic encoding during decision-making. Nature neuroscience 23, 1410–1420. 10.1038/s41593-020-0696-5.

46. Sarel, A., Palgi, S., Blum, D., Aljadeff, J., Las, L., and Ulanovsky, N. (2022). Natural switches in behaviour rapidly modulate hippocampal coding. Nature 609, 119–127. 10.1038/s41586-022-05112-2.

47. Kim, J., Joshi, A., Frank, L., and Ganguly, K. (2023). Cortical-hippocampal coupling during manifold exploration in motor cortex. Nature 613, 103–110. 10.1038/s41586-022-05533-z.

48. Steinmetz, N.A., Zatka-Haas, P., Carandini, M., and Harris, K.D. (2019). Distributed coding of choice, action and engagement across the mouse brain. Nature 576, 266-+. 10.1038/s41586-019-1787-x.

49. Allen, W.E., Chen, M.Z., Pichamoorthy, N., Tien, R.H., Pachitariu, M., Luo, L.Q., and Deisseroth, K. (2019). Thirst regulates motivated behavior through modulation of brainwide neural population dynamics. Science 364, 253-+. 10.1126/science.aav3932.

50. Kennedy, A., Kunwar, P.S., Li, L.Y., Stagkourakis, S., Wagenaar, D.A., and Anderson, D.J. (2020). Stimulus-specific hypothalamic encoding of a persistent defensive state. Nature 586, 730–734. 10.1038/s41586-020-2728-4.

51. Yang, T., Yang, C.F., Chizari, M.D., Maheswaranathan, N., Burke, K.J., Borius, M., Inoue, S., Chiang, M.C., Bender, K.J., Ganguli, S., and Shah, N.M. (2017). Social Control of Hypothalamus-Mediated Male Aggression. Neuron 95, 955-+. 10.1016/j.neuron.2017.06.046.

52. Ishii, K.K., Osakada, T., Mori, H., Miyasaka, N., Yoshihara, Y., Miyamichi, K., and Touhara, K. (2017). A Labeled-Line Neural Circuit for Pheromone-Mediated Sexual Behaviors in Mice. Neuron 95, 123-+. 10.1016/j.neuron.2017.05.038.

53. Falkner, A.L., Grosenick, L., Davidson, T.J., Deisseroth, K., and Lin, D.Y. (2016). Hypothalamic control of male aggression-seeking behavior. Nature neuroscience 19, 596-+. 10.1038/nn.4264.

54. Lee, H., Kim, D.W., Remedios, R., Anthony, T.E., Chang, A., Madisen, L., Zeng, H.K., and Anderson, D.J. (2014). Scalable control of mounting and attack by Esr1+ neurons in the ventromedial hypothalamus. Nature 509, 627-+. 10.1038/nature13169.

55. Yang, C.F., Chiang, M.C., Gray, D.C., Prabhakaran, M., Alvarado, M., Juntti, S.A., Unger, E.K., Wells, J.A., and Shah, N.M. (2013). Sexually Dimorphic Neurons in the Ventromedial Hypothalamus Govern Mating in Both Sexes and Aggression in Males. Cell 153, 896–909. 10.1016/j.cell.2013.04.017.

56. Liu, D., Rahman, M., Johnson, A., Tsutsui-Kimura, I., Pena, N., Talay, M., Logeman, B.L., Finkbeiner, S., Choi, S., Capo-Battaglia, A., et al. (2023). A Hypothalamic Circuit Underlying the Dynamic Control of Social Homeostasis. bioRxiv. 10.1101/2023.05.19.540391.

57. Stagkourakis, S., Smiley, K.O., Williams, P., Kakadellis, S., Ziegler, K., Bakker, J., Brown, R.S.E., Harkany, T., Grattan, D.R., and Broberger, C. (2020). A Neuro-hormonal Circuit for Paternal Behavior Controlled by a Hypothalamic Network Oscillation. Cell 182, 960-+. 10.1016/j.cell.2020.07.007.

58. Mei, L., Yan, R.Z., Yin, L.P., Sullivan, R.M., and Lin, D.Y. (2023). Antagonistic circuits mediating infanticide and maternal care in female mice. Nature. 10.1038/s41586-023-06147-9.

59. Zhang, S.X., Lutas, A., Yang, S., Diaz, A., Fluhr, H., Nagel, G., Gao, S.Q., and Andermann, M.L. (2021). Hypothalamic dopamine neurons motivate mating through persistent cAMP signalling. Nature 597, 245-+. 10.1038/s41586-021-03845-0.

60. Kohl, J., and Dulac, C. (2018). Neural control of parental behaviors. Current opinion in neurobiology 49, 116–122. 10.1016/j.conb.2018.02.002.

61. Kira, S., Safaai, H., Morcos, A.S., Panzeri, S., and Harvey, C.D. (2023). A distributed and efficient population code of mixed selectivity neurons for flexible navigation decisions. Nature communications 14, 2121. 10.1038/s41467-023-37804-2.

62. Kaufman, M.T., Benna, M.K., Rigotti, M., Stefanini, F., Fusi, S., and Churchland, A.K. (2022). The implications of categorical and category-free mixed selectivity on representational geometries. Current opinion in neurobiology 77, 102644. 10.1016/j.conb.2022.102644.

63. Johnston, W.J., Palmer, S.E., and Freedman, D.J. (2020). Nonlinear mixed selectivity supports reliable neural computation. PLoS computational biology 16, e1007544. 10.1371/journal.pcbi.1007544.

64. Grunfeld, I.S., and Likhtik, E. (2018). Mixed selectivity encoding and action selection in the prefrontal cortex during threat assessment. Current opinion in neurobiology 49, 108–115. 10.1016/j.conb.2018.01.008.

65. Rigotti, M., Barak, O., Warden, M.R., Wang, X.J., Daw, N.D., Miller, E.K., and Fusi, S. (2013). The importance of mixed selectivity in complex cognitive tasks. Nature 497, 585–590. 10.1038/nature12160.

66. Schoonover, C.E., Ohashi, S.N., Axel, R., and Fink, A.J.P. (2021). Representational drift in primary olfactory cortex. Nature 594, 541–546. 10.1038/s41586-021-03628-7.

67. Zhao, P., Chen, X., Bellafard, A., Murugesan, A., Quan, J., Aharoni, D., and Golshani, P. (2023). Accelerated social representational drift in the nucleus accumbens in a model of autism. bioRxiv. 10.1101/2023.08.05.552133.

68. Haimerl, C., Savin, C., and Simoncelli, E.P. (2019). Flexible information routing in neural populations through stochastic comodulation. Advances in Neural Information Processing Systems 32 (Nips 2019) 32.

69. Guo, Z., Yin, L., Diaz, V., Dai, B., Osakada, T., Lischinsky, J.E., Chien, J., Yamaguchi, T., Urtecho, A., Tong, X., et al. (2023). Neural dynamics in the limbic system during male social behaviors. Neuron. 10.1016/j.neuron.2023.07.011.

70. Segalin, C., Williams, J., Karigo, T., Hui, M., Zelikowsky, M., Sun, J.J., Perona, P., Anderson, D.J., and Kennedy, A. (2021). The Mouse Action Recognition System (MARS) software pipeline for automated analysis of social behaviors in mice. eLife 10. 10.7554/eLife.63720.

71. Jun, J.J., Steinmetz, N.A., Siegle, J.H., Denman, D.J., Bauza, M., Barbarits, B., Lee, A.K., Anastassiou, C.A., Andrei, A., Aydin, C., et al. (2017). Fully integrated silicon probes for high-density recording of neural activity. Nature 551, 232-+. 10.1038/nature24636.

72. Lundberg, S.M., and Lee, S.I. (2017). A Unified Approach to Interpreting Model Predictions. Adv Neur In 30.

73. Chen, T.Q., and Guestrin, C. (2016). XGBoost: A Scalable Tree Boosting System. Kdd’16: Proceedings of the 22nd Acm Sigkdd International Conference on Knowledge Discovery and Data Mining, 785–794. 10.1145/2939672.2939785.

